# The zinc finger proteins ZC3H20 and ZC3H21 stabilise mRNAs encoding membrane proteins and mitochondrial proteins in insect-form *Trypanosoma brucei*

**DOI:** 10.1101/646547

**Authors:** Bin Liu, Kevin Kamanyi Marucha, Christine Clayton

## Abstract

ZC3H20 and ZC3H21 are related trypanosome proteins with two C(x)_8_C(x)_5_C(x)_3_H zinc finger motifs. ZC3H20 is unstable in mammalian-infective bloodstream forms, but becomes more abundant as they transform to growth-arrested stumpy form, while ZC3H21 appears only in the procyclic form of the parasite, which infects Tsetse flies. Each protein binds to several hundred mRNAs, with overlapping but not identical specificities. Both increase expression of bound mRNAs, probably through recruitment of the MKT1-PBP1 complex. At least seventy of the bound mRNAs decrease after RNAi targeting ZC3H20 or ZC3H20 and ZC3H21; their products include procyclic-specific proteins of the plasma membrane and energy metabolism. Simultaneous depletion of ZC3H20 and ZC3H21 causes procyclic forms to shrink and stop growing; in addition to decreases in target mRNAs, there are other changes suggestive of loss of developmental regulation. The bloodstream-form specific protein RBP10 controls ZC3H20 and ZC3H21 expression. Interestingly, some ZC3H20/21 target mRNAs also bind to and are repressed by RBP10, allowing for dynamic regulation as RBP10 decreases and ZC3H20 and ZC3H21 increase during differentiation.

## Introduction

The unicellular eukaryote *Trypanosoma brucei* is a useful model organism for studies of post-transcriptional regulation of gene expression. As in other Kinetoplastids, transcription in *T. brucei* is polycistronic, and mRNAs are excised by co-transcriptional *trans* splicing and polyadenylation. RNA binding proteins play important roles throughout the lifetime of every mRNA, governing mRNA processing, translation, and mRNA decay (Clayton, 2019, Kolev *et al.*, 2014).

*T. brucei* is always extracellular, and grows in mammals as long slender trypomastigotes. These have a variant surface glycoprotein surface coat and depend mainly on glycolysis for ATP, using enzymes that are compartmentalised in a microbody called the glycosome (Hannaert *et al.*, 2003). When bloodstream forms reach high density, either in culture or in a mammal, they have a quorum sensing response (Rojas *et al.*, 2018) and transform to non-dividing stumpy-form trypomastigotes. Upon uptake of the stumpy forms by Tsetse, or transfer to culture at 27°C in the presence of cis-aconitate, stumpy forms develop into growing procyclic trypomastigotes, which rely mainly on mitochondrial pathways of ATP generation (Mantilla *et al.*, 2017, Bringaud *et al.*, 2006). The trypanosomes later migrate via the proventriculus to the salivary glands, becoming first epimastigotes, then non-dividing metacyclic trypomastigotes which have variant surface glycoprotein and are infectious for mammals (Rotureau & Van Den Abbeele, 2013). All of these transitions are accompanied by changes in RNA-binding proteins, and in some cases are known to be governed by them (Kolev *et al.*, 2012, Kolev *et al.*, 2014, Mugo & Clayton, 2017).

One example of a controlling RNA binding protein is ZC3H11, which has a single CCCH zinc finger domain (consensus: C(x)_8_C(x)_5_C(x)_3_H) and promotes the expression of chaperone mRNAs. ZC3H11 binds (UAA) repeats in mRNA 3’-untranslated regions (3’-UTRs), and is required for stabilisation of its target mRNAs after heat shock (Droll *et al.*, 2013). ZC3H11 recruits another protein, MKT1, via an interaction motif, (Y/W/V/I)(R/T/Q)H(N/D)PY. MKT1 is part of a protein complex that promotes mRNA translation and stability (Singh *et al.*, 2014).

A second RNA-binding protein, RBP10, is essential to maintain the differentiation state of bloodstream form *T. brucei*. RBP10 binds to the sequence UA(U)_6_ in 3’-UTRs, targeting the mRNAs for translational repression and destruction (Mugo & Clayton, 2017). Levels of RBP10 decline during stumpy formation. In procyclic forms, RBP10 is undetectable and its target mRNAs are generally abundant and well translated. Correspondingly, depletion of RBP10 in bloodstream forms triggers a cascade of events, starting with increases in RBP10 target mRNAs, and followed by secondary effects that switch the gene expression pattern towards that of procyclic forms. Conversely, expression of RBP10 in procyclic forms switches the cells towards the bloodstream-form pattern (Mugo & Clayton, 2017).

Among the directly bound targets of RBP10 in bloodstream forms are the mRNAs encoding three proteins with CCCH zinc finger domains: ZC3H22 (Tb927.7.2680), ZC3H21 (Tb927.7.2670), and ZC3H20 (Tb927.7.2660). The 3’-UTRs of these mRNAs have, respectively, two, five, and seven copies of the RBP10 consensus recognition motif (Fig 1A). All three mRNAs are more abundant, and considerably better translated, in procyclic forms than in growing bloodstream forms (Fadda *et al.*, 2014, Dejung *et al.*, 2016, Jensen *et al.*, 2014, Antwi *et al.*, 2016), and their expression is also lower in metacyclic forms than in procyclic forms (Christiano *et al.*, 2017). When ZC3H20 or ZC3H21 are tethered (via a lambdaN-peptide-boxB interaction) to a reporter mRNA in bloodstream forms, expression is stimulated, whereas tethering of ZC3H22 leads to repression (Lueong *et al.*, 2016, Erben *et al.*, 2014). Tagged versions of each are in the cytoplasm of procyclic forms (see www.tryptag.org) (Dyer *et al.*, 2016), and move to messenger ribonucleoprotein (mRNP) granules after starvation (Fritz *et al.*, 2015).

**Fig 1.**
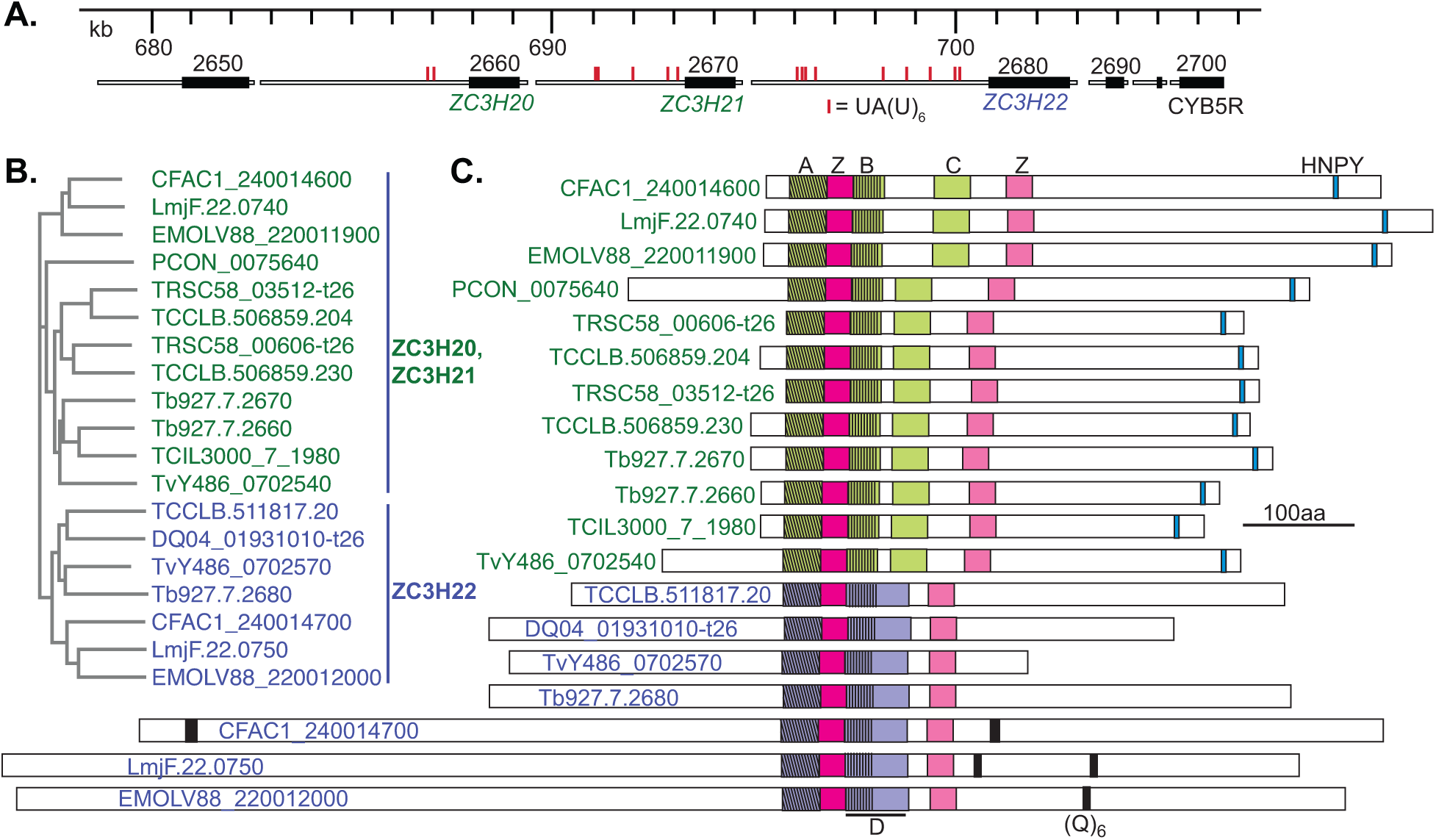
ZC3H20,21 and 22 in different Kinetoplastids. A) Map of the Trypanosoma brucei TREU927 ZC3H20, ZC3H21 and ZC3H22 genes, taken from TritrypDB. Open reading frames are in black. The narrower grey untranslated regions (UTRs) have been modified from the database annotation to reflect the long 3’-UTRs seen by RNASeq and Northern blotting. B) Tree of full-length proteins made with Clustal Omega (https://www.ebi.ac.uk/Tools/msa/clustalo/), showing real branch lengths. ZC3H20/21 are in green and ZC3H22 in blue. Organism abbreviations are: CFAC: *Crithidia fasciculata*; Lmj: *Leishmania major*; EMOLV: *Endotrypanum monterogeii*; PCON: *Paratrypanosoma confusum*; TRSC: *Trypanosoma rangeli*; TCCLB: *Trypanosoma cruzi*; Tb: *Trypanosoma brucei brucei;* TCIL: *Trypanosoma congolense;* Tv: *Trypanosoma vivax.* To obtain the maps, *T. congolense* and *T. vivax* complete open reading frames were reconstructed by deleting individual nucleotides from the sequence assemblies. TRSC58_03512-t26 C) Maps of the proteins showing conserved regions. The first CCCH domain is in dark pink and the second one is in pale pink (both marked “Z” at the top of the diagram. Regions conserved in ZC3H20 homologues are in green, and those conserved in ZC3H22 homologues are in purple. The cross-hatching indicates regions that are partially conserved between ZC3H21 and 22. Conservation was as follows: A: 15/33 identical, 20/33 similar within ZC3H21 homologues, 19/33 identical, 21/33 similar within ZC3H22 homologues, and 12/33 identical in all proteins shown; B: 13/27 identical within ZC3H20 homologues, 8/24 identical in all proteins shown; C: 5/33 identical, 11/33 similar within ZC3H20 homologues; D: 19/62 identical, 24/62 similar within ZC3H22 homologues. HNPY-related motifs are indicated in cyan and polyglutamine tracts (at least 6 glutamines, (Q)_6_) in black. An alignment of the zinc-finger-domaincontaining region is provided as supplementary Fig S1. Additional possible divergent CCCH sequences towards the C-termini of ZC3H20 and ZC3H21 are not conserved and are not shown. They are C(X)_5_C(X)_8_C(X)_3_H in ZC3H20 and C(X)_12_C(X)_5_C(X)_3_H in ZC3H21.

Ling et al (Ling *et al.*, 2011) previously found that RNAi that simultaneously targeted *ZC3H20* and *ZC3H21* inhibited procyclic-form trypanosome growth. Microarray experiments revealed that RNAi resulted in reduced levels of six RNAs, and increases in another ten. The authors also demonstrated binding of ZC3H20 to two of the reduced transcripts. To obtain a detailed picture of the functions of ZC3H20 and ZC3H21 we have now determined their transcriptome-wide binding specificities. These, combined with analyses of expression and the effects of depletion, showed us that the two proteins affect the fates of several hundred mRNAs, and have overlapping but different roles in the trypanosome life cycle and in gene regulation.

## Results

### Conservation of ZC3H20, 21 and 22

The ZC3H22, ZC3H21 and ZC3H20 genes are arranged consecutively in the *Trypanosoma brucei* genome (Fig 1A). ZC3H21 and ZC3H20 are similar in the region that includes the two zinc fingers, each of which has the structure C(x)_7_C(x)_5_C(x)_3_H (Fig 1B, C and Supplementary Fig S1). ZC3H22 is similar to the other two proteins only in the region containing the N-terminal zinc finger; its C-terminal one has the structure C(x)_8_C(x)_5_C(x)_3_H. There is at least one homologue of ZC3H20 or ZC3H21, and of ZC3H22, in *Trypanosoma, Leishmania, Crithidia, Endotrypanum, Leptomonas, Paratrypanosoma and Blechomonas* species (Fig 1B, C Supplementary Fig S1) but both ZC3H21 and ZC3H22 appear to be absent in the free-living *Bodo caudatus*. In *Trypanosoma congolense* and *Trypanosoma vivax*, for which the genomes are incomplete, all three genes appear to be present only as fragments, but longer open reading frames are obtained by adding or deleting a few nucleotides, yielding homologues containing both zinc fingers, which suggests that the fragmentation might be due to assembly errors. The phylogenetic tree (Fig 1B) suggests that duplication of ZC3H21 occurred independently in *Trypanosoma bruce*i and in *T. cruzi* (TCCLB) and *T. rangeli* (TRSC), which are transmitted by triatomine bugs. The duplication may therefore be an adaptation to specific conditions.

Apart from the conservation around the zinc fingers, all ZC3H20 and ZC3H21 sequences have a variant of the MKT1-binding motif (Y/W/T)(R/T/Q)H(N/D)PY near their C-termini. The C-termini of the different ZC3H22s, in contrast, have little in common. Three of the ZC3H22s do, however, have poly(Q) sequences that might form disordered interaction domains (Fig 1C, Supplementary Fig S1).

The RNA-binding specificities of CCCH domains are influenced by the sequences within the zinc fingers, and by 5 flanking residues (Hudson *et al.*, 2004, Pagano *et al.*, 2007). The N-terminal CCCH domains of ZC3H20 and ZC3H21 have only three differences, but the C-terminal motifs have eight (Supplementary Fig S2). This suggests that the RNA-binding specificities of the two proteins could be related, but not identical. ZC3H22 specificity is likely to differ more. A comparison with the RNA-binding regions of other *T. brucei* proteins with two CCCH domains revealed that the each is likely to have binding specificities that differ from those of ZC3H20 and ZC3H21 (Supplementary Fig S2). The proteins with most closely related zinc finger domains are ZC3H18 and ZC3H30.

### Regulated expression of ZC3H20 and 21

Unless otherwise stated, experiments in bloodstream forms were done with differentiation-competent EATRO1125 strain trypanosomes. They were grown in methyl-cellulose-containing medium (Vassella *et al.*, 2001) and their density was routinely maintained below 8×10^5^ cells/ml, in order to maintain differentiation competence. Some experiments were also done with monomorphic Lister 427 strain bloodstream-form *T. brucei*, which cannot make stumpy forms (Vassella *et al.*, 1997). Established procyclic-form cultures were either Lister 427 or EATRO1125 strain trypanosomes.

We first confirmed available data concerning developmental regulation of ZC3H20 and ZC3H21, since mass spectrometry results on total lysates have limited sensitivity. The 5’-UTRs of *T. brucei* genes have not so far been implicated in developmental regulation. Protein expression can therefore be assessed using N-terminally *in-situ*-tagged versions, with the caveat that N-terminal tagging might influence protein stability. For maximal sensitivity, we used versions bearing a tandem affinity purification (TAP) tag that includes an IgG binding domain, allowing detection by direct IgG binding (Puig *et al.*, 2001). Functionality of the tagged protein was confirmed by deletion of the remaining wild-type allele (Fig 2B, Supplementary Fig S3 B, H) but for the differentiation analysis we used cells that also had wild-type protein. TAP-ZC3H20 was just detectable in long slender differentiation-competent bloodstream-form trypanosomes. As predicted by proteomic results (Dejung *et al.*, 2016), TAP-ZC3H20 increased at maximal density and was more abundant in trypanosomes that had been further incubated at high density at 37°C (Fig 3A, B and Supplemental Fig S4A). The increase in ZC3H20 preceded detection of the stumpy-form marker PAD1 (Dean *et al.*, 2009) (Fig 3B).

**Fig. 2.**
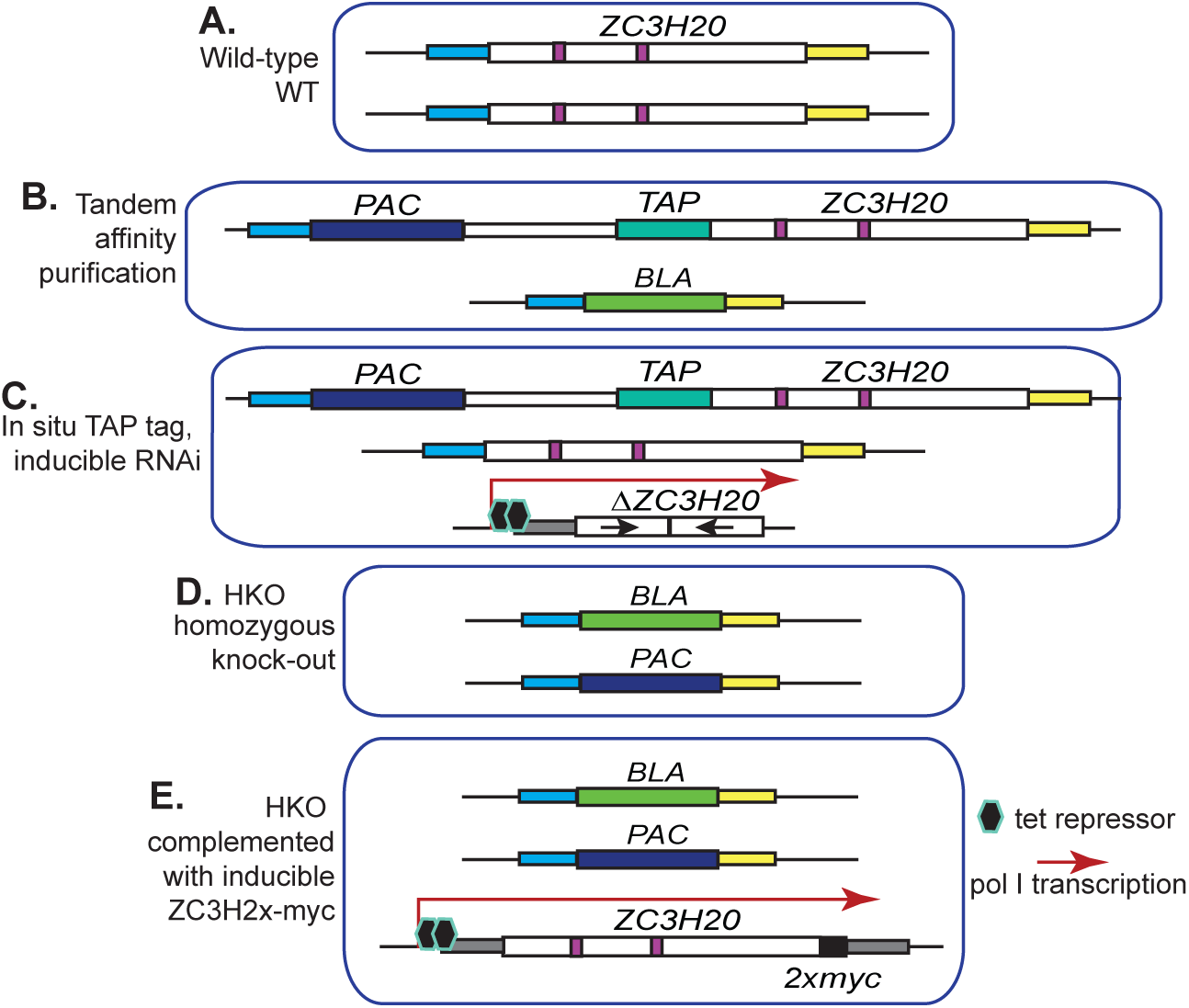
Genotypes of transgenic cell lines with altered ZC3H20 expression. The different panels are explained in the text and on the Figure. Additional genotypes and examples of PCR evidence are illustrated in Supplementary Fig S3.

**Fig 3.**
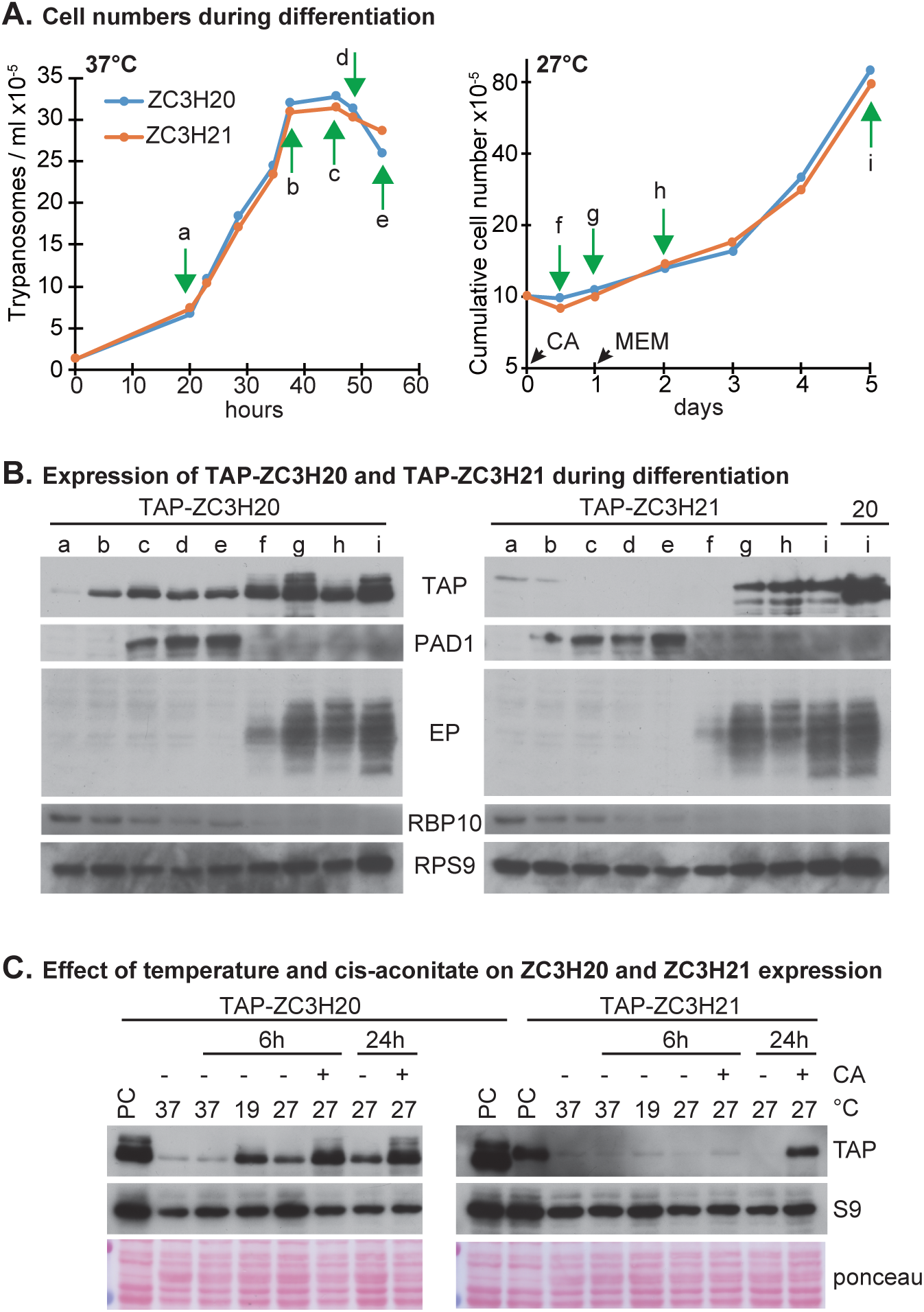
Expression of ZC3H20 and ZC3H21. A. Cell numbers during differentiation of bloodstream forms. EATRO1125 bloodstream forms expressing N-terminally TAP-tagged ZC3H20 (Z20) or ZC3H21 (Z21) were used (Supplementary Fig S3B, H). Initial growth (left panel) was in HMI9 at 37°C with methylcellulose. At time point d, cis-aconitate was added to trigger differentiation and the temperature was lowered to 27°C. 1 day later the cells were transferred to MEM-Pros medium (right panel). Cells were harvested at different time points (a-i) for Western blot analysis. B. Western blots for the experiment in panel (A). The proteins detected are indicated next to the panels. On the far right of the TAP-ZC3H21-TAP blot, the same number of cells expressing TAP-ZC3H20 have been loaded. C. Effect of temperature reduction on expression of TAP-ZC3H20 and TAP-ZC3H21. Cells expressing the tagged proteins and grown initially at 37°C were incubated at a starting density of 2.5×10^5^/ml, for the times indicated, at lower temperatures. CA: cis-aconitate.

To induce differentiation of stumpy forms to procyclic forms (Fig 3A), the temperature was decreased from 37°C to 27°C, and cis-aconitate (6mM) was added. 24h later the cells were transferred to procyclic-form medium, which has proline as the principal energy source. Growth resumed within 24h. Again as predicted from proteomics, ZC3H20 expression increased only slightly then remained constant (Fig 3A, B, Supplemental Fig S4 A). Interestingly, TAP-ZC3H20 also increased when monomorphic Lister 427 strain bloodstream forms attained high density (Supplemental Fig S4 B). This suggests that the increase in TAP-ZC3H20 does not depend the ability to make stumpy forms in response to the quorum-sensing signal (Vassella *et al.*, 1997). The increase in TAP-ZC3H20 correlated with the stationary phase of culture and was not dependent on the presence of methyl cellulose (Supplemental Fig S4 B, C).

TAP-ZC3H21 was, like TAP-ZC3H20, just detectable in bloodstream forms, but it did not increase upon incubation at high density (Fig 3A, B, Supplemental Fig S4D), which is consistent with its absence in stumpy-form proteomes (Dejung *et al.*, 2016). TAP-ZC3H21 only became readily detectable upon induction of procyclic differentiation, and it was not as abundant as TAP-ZC3H20 in established procyclic forms (Fig 3A, B, Supplemental Fig S4D).

We next examined which differentiation stimuli affected ZC3H20 and ZC3H21 expression. Treatment of growing (low density) EATRO1125 bloodstream forms at 27°C alone caused an increase in ZC3H20 (Fig 3C, Supplementary Fig S4E). Incubation at 19°C sensitizes bloodstream forms to differentiation signals (Engstler & Boshart, 2004). This caused a mild increase in ZC3H20, but higher expression was obtained at 27°C in the presence of cis-aconitate (Fig 3C, Supplementary Fig S4G). An increase in TAP-ZC3H21 at 19°C was seen once (Supplementary Fig S4H), but it was only reproducibly increased after both 27°C and cis-aconitate treatment (Fig 3C, Supplementary Fig S4H)., Incubation of procyclic forms at high density did not affect TAP-ZC3H20 expression (Supplementary Fig S4F).

The *ZC3H20* and *ZC3H21* mRNAs were not examined separately in our previous transcriptome analyses (Mugo & Clayton, 2017). The *ZC3H20* 3’-UTR has two RBP10-binding motifs, whereas that of *ZC3H21* has five. A pull-down confirmed that *ZC3H20* mRNA is bound by RBP10 (Fig 4A). TAP-ZC3H20 expression in bloodstream forms increased after *rbp10* RNAi, as expected (Fig 4B).

**Fig 4.**
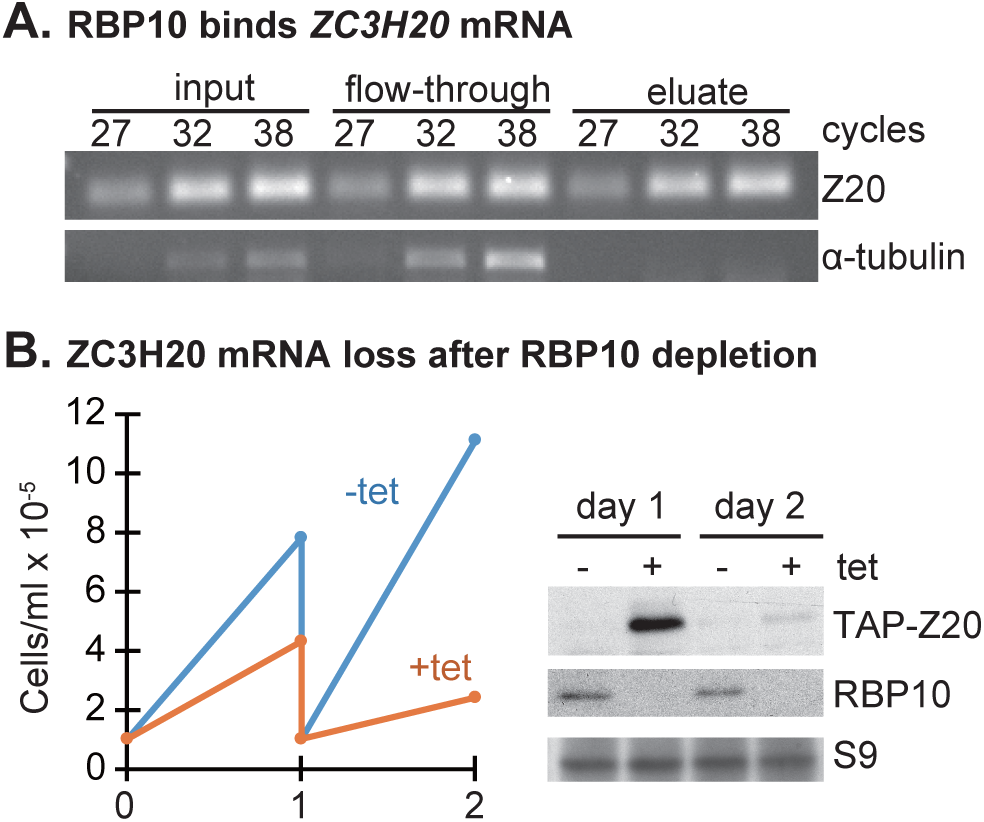
ZC3H20 expression control by RBP10. A. Immunoprecipitation of TAP-RBP10 was performed in bloodstream form Lister 427 *T. brucei*. RNA was purified from the input, flowthrough and eluate fractions. ZC3H20 and tubulin (negative control) mRNAs were detected using RT-PCR. B. *RBP10* was targeted by RNAi in Lister 427 bloodstream form T. brucei expressing TAPZC3H20. Cell densities are shown on the left and expression of ZC3H20 (Z20) and RBP10 on the right.

### Activation by tethered ZC3H20 and ZC3H21

In tethering screens in bloodstream forms, full-length λN-ZC3H20 increased reporter expression, as did a C-terminal fragment starting at residue 222 (Lueong *et al.*, 2016, Erben *et al.*, 2014). To confirm this for procyclic forms, we expressed λN-ZC3H20-myc in cells containing a *CAT*-boxB mRNA that has five boxB sequences between the *CAT* open reading frame and the actin 3’-UTR (Fig 5A). A reporter lacking boxB served as the control. Tethering of the full-length protein (Fig 5B) gave a 2-4 fold increase in CAT activity (Fig 5C) and also increased the RNA (Fig 5D).

**Fig. 5.**
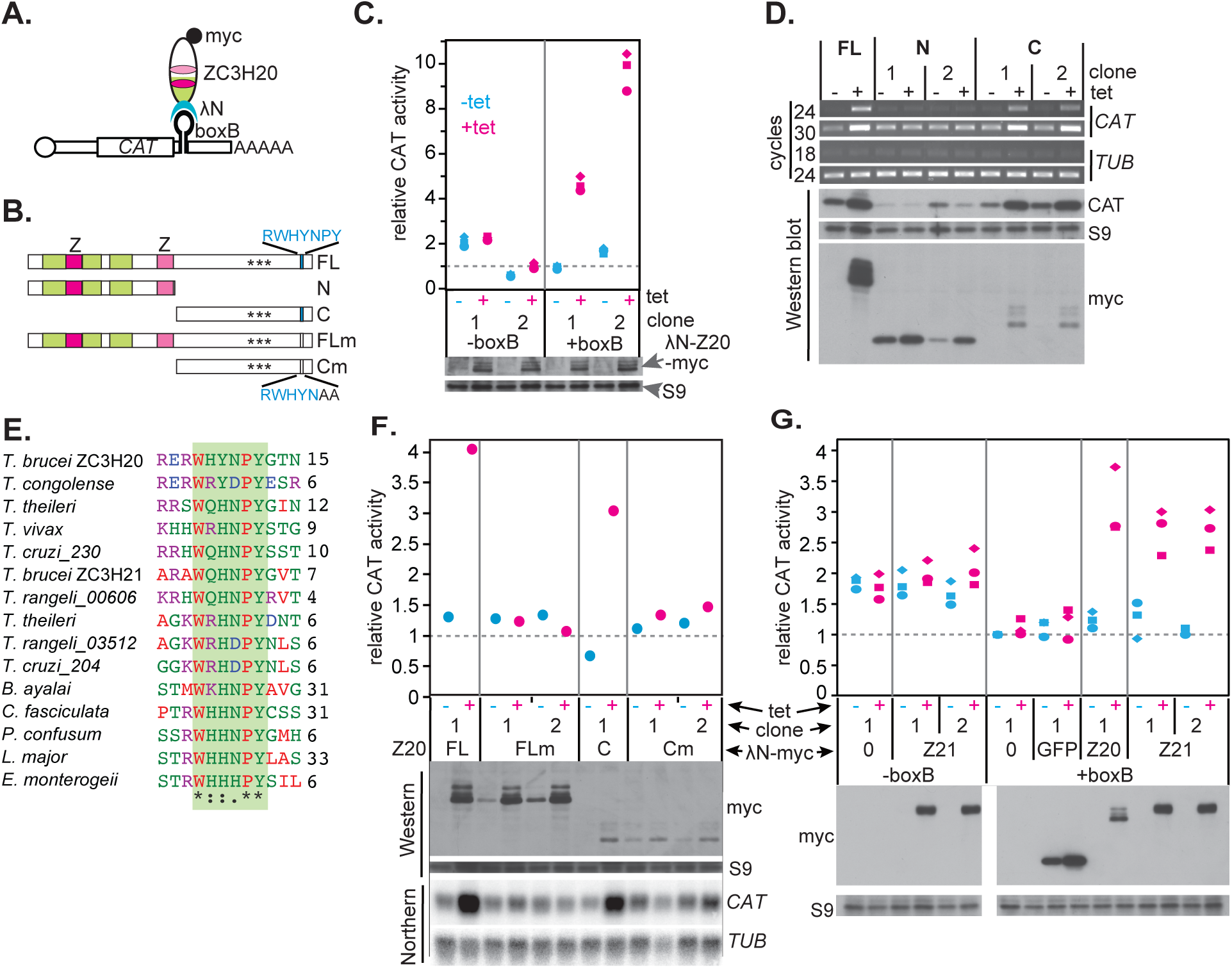
Enhancement of expression by tethered ZC3H20 in procyclic forms depends on the MKT1 interaction motif. A. The cartoon shows λN-ZC3H20-myc bound to a boxB motif in the *CAT*-boxB mRNA, which has five boxB sequences between the *CAT* open reading frame and the actin 3’-UTR. B. Cartoon of ZC3H20 fragments used in the tethering assay. Colour codes are as in Fig 1. FL is full-length protein, “N” is the N-terminal part of the protein containing the RNA binding motifs, and “C” is the C-terminal part of the protein containing some phosphorylation sites and the putative MKT1 interacting motif. The lowest diagrams show full-length or C-terminal portions with point mutations in the motif. C. Relative CAT (chloramphenicol acetyltransferase) activities in cells expressing reporters with or without boxB (two clones each, C1 and C2). Expression of λN-ZC3H20-myc was assessed by western blot (lower panel). D. Effect of tethering λN-ZC3H20-myc fragments. Upper four panels: Levels of *CAT* mRNA were assessed by detected by RT-PCR with two different numbers of cycles and with tubulin as the control. Lower panels: Expression of CAT and of the λN-ZC3H20-myc proteins, assessed by Western blotting with ribosomal protein S9 as the control. E. Conservation of the MKT1-interacting motif in ZC3H20/21 homologues and paralogues. The three amino acids flanking either side of the motif are included, and the numbers of amino acids to the C-termini are shown. Full protein IDs are in supplementary Fig S1. F. Effect of mutating the MKT1 interaction motif. Different λN-ZC3H20-myc proteins (see panel B) were used. The graph shows CAT activity, and beneath are Northern and Western blots. G. Tethering of λN-ZC3H21-myc.

ZC3H20 and ZC3H21 interacted with MKT1 in the yeast 2-hybrid assay (Singh *et al.*, 2014) and all ZC3H20/ZC3H21 homologues contain a variant of the MKT1-binding motif, (Y/W/T)(R/T/Q)H(N/D)PY (Fig 5E). However, the motif in *T. brucei* ZC3H20, WHYNPY, which starts at position 383, does not conform to the consensus because it has an inserted tyrosine residue. To find sequences required for activation by tethered ZC3H20 we first tethered N-and C-terminal λN-ZC3H20-myc fragments (Fig 5B). The results (Fig 5F) confirmed that activation was mediated by the C-terminal half of the protein. Next, we mutated the most conserved residues of the motif, proline-tyrosine (Fig 5B, E), to alanines (Fig 5B). This mutation was previously shown to eliminate the interaction between ZC3H11 and MKT1 (Singh *et al.*, 2014). The mutated protein indeed lacked activity (Fig 5F). We concluded that activation by ZC3H20 in the tethering assay probably depends on recruitment of MKT1. As expected, λN-ZC3H21-myc, which has C-terminal WQHNPY, also had an activating effect (Fig 5G).

### ZC3H20 is required during differentiation to the stumpy and procyclic forms

To assess the requirement for ZC3H20 in differentiation we first attempted depletion by RNAi. We used cell lines expressing TAP-ZC3H20 in order to be able to follow protein loss (Fig 2C). Two independent EATRO1125 clones were grown to 3×10^6^/ml in methyl cellulose medium and incubated for a further 12 h, at which point PAD1 was expressed (Fig 6A). Tetracycline was then added and differentiation was induced without methyl cellulose, by addition of cis-aconitate and reduction of the temperature to 27°C. One day later, cells were transferred to procyclic medium. Induction of dsRNA transcription was initially poor, as judged by a rather small decrease in TAP-ZC3H20 relative to the control; this is probably because the cells were approaching stationary phase. Nevertheless, after 1 day, expression of the procyclic surface protein EP procyclin was impaired in the induced lines (Fig 6A, upward arrows). This suggested that ZC3H20 was required for normal differentiation kinetics. After 5 days, the cells were growing normally as procyclic forms, but some ZC3H20 was still detectable, so its role could not be assessed.

**Fig 6.**
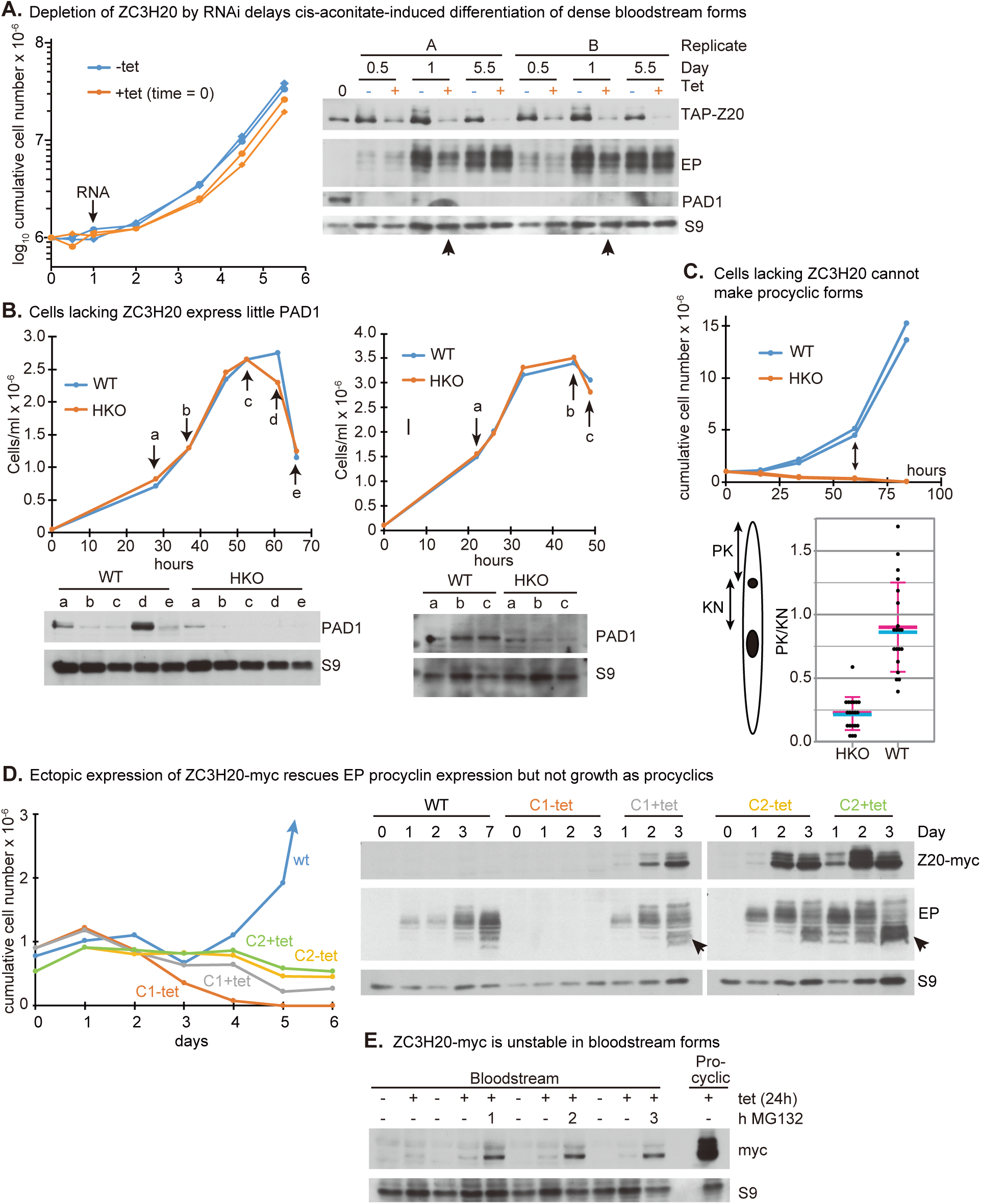
Depletion of ZC3H20 affects differentiation. A. The cells used were EATRO1125 bloodstream forms expressing TAP-ZC3H20 and with tetracycline-inducible RNAi targeting *ZC3H20* (Fig 2C). The cells were maintained at high density with methyl cellulose (around 3×10^6^/mL) for 12 hours prior to the start of the experiment. At time=0, cells were transferred to medium without methyl cellulose (1×10^6^/mL), cis-aconitate was added with or without 100ng/ml tetracycline, and the temperature was reduced to 27°C. One day later cells were switched to MEM-pros medium. The experiment was done in two independent cultures (A and B). Cell densities for the two clones (with different symbols) are shown on the left; the arrow indicates the time at which some of the cells were harvested for RNASeq. Western blots for protein expression on the right; the arrows point to RNAi lanes with lower EP procyclin expression than the controls. S9 is ribosomal protein S9. B. Bloodstream-form EATRO1125 *T. brucei* that lack ZC3H20 (Fig 2D, Supplementary Fig S3G) do not express PAD1 at high density. Cells were allowed to grow in methyl cellulose without dilution. The cell densities are shown above and Western blots beneath. The antibody to PAD1 cross-reacts with other proteins, which may explain the band in HKO at time point “a”. Results from two different experiments are shown. C. Cells lacking ZC3H20 (HKO, Fig 2D) were grown to 3×10^6^/ml, then treated as in (A) except that the cells were maintained at high density for 16h before cis-aconitate addition at 1x 10^6^/ml. Cells with two *ZC3H20* genes served as controls. The upper panel shows cumulative cell numbers - the cells lacking ZC3H20 died. The lower panel illustrates the morphologies of the cells after 60h (double-headed arrow). P=posterior, K= kinetoplast, N=nucleus. A higher PK/KN ratio is characteristic of procyclic forms. Cyan: median; magenta: arithmetic mean and standard deviation. D. Inducible ZC3H20-myc only partially complements the lack of ZC3H20. EATRO1125 *T. brucei* were used in which lacked *ZC3H20* genes (HKO) but had a tetracycline-inducible copy of ZC3H20-myc (Fig 2E). Two clones, C1 and C2, were compared and wild-type was used as a control. At time point 0, long slender bloodstream-form cells were treated with cis-aconitate with or without tetracycline. Cumulative cell numbers are on the left and protein expression on the right. E. Exponentially growing bloodstream forms with inducible ZC3H20-myc (clone 1 from panel D) were incubated with or without tetracycline for 24h, then MG132 was added for the indicated times. Procyclic forms with inducible lambdaN-ZC3H20-myc, also induced for 24h, are shown for comparison.

To further investigate ZC3H20 function, we deleted both genes in bloodstream forms (homozygous knock-out, HKO) (Fig 2D, Supplementary Fig S3 E and G). These cells were unable to express the stumpy-form marker PAD1 at high density (Fig 6B). When procyclic-form differentiation was induced the cells were unable to proliferate, failed to switch their morphology and eventually died (Fig 6C). To check whether the defect was indeed due to loss of ZC3H20, we introduced a construct enabling tetracycline-inducible expression of C-terminally myc-tagged ZC3H20 (ZC3H20-myc) into the HKO bloodstream forms (Fig 2E). Preliminary results suggested that the protein was not expressed after tetracycline addition, but nevertheless, we attempted differentiation. Since there is little transcription in stumpy forms, we used growing cells; tetracycline and cis-aconitate were added and the temperature was reduced to 27°C. After a further day, the medium was changed as before. With this protocol, wild-type cells start to grow as procyclic forms on day 4, with detectable EP procyclin expression after 1 day (Fig 6D). For inducible ZC3H20-myc, two different cloned lines were studied (Fig 6D and supplementary Fig S6A). In the absence of tetracycline, clone 1 showed no expression of ZC3H20-myc and was dying by day 3. With tetracycline, ZC3H20-myc appeared gradually over three days, procyclin expression was restored, and survival of the cells was prolonged - but they never grew as procyclic forms (Fig 6D, supplementary Fig S6A). This suggested that ZC3H20-myc was functional in the early stages of differentiation. Clone 2 showed expression of ZC3H20 both with and without tetracycline, and behaved similarly to clone 1 in the presence of tetracycline. In both cases, however, EP procyclin expression pattern was sometimes altered, with accumulation on day 3 of faster-migrating proteins that were present only at low abundance in the wild-type cells (Fig 6D and Supplementary Fig S6A, arrows). Either post-translational modification of the protein was defective, or different procyclin genes were being expressed.

These results suggested that excess ZC3H20 might prevent establishment of growing procyclic forms. We therefore examined differentiation-competent bloodstream-form trypanosomes with inducibly expressed ZC3H20-myc as well as normal *ZC3H20* genes (Supplementary Fig S3K), using the same protocol as before. Again, addition of tetracycline to the growing bloodstream forms gave no detectable ZC3H20-myc expression, and had no effect on growth (not shown). In the absence of tetracycline, one clone had no detectable ZC3H20-myc and could differentiate and grow as procyclic forms, but this was prevented by tetracycline addition. Another clone had less regulation and was unable to grow as procyclic forms (Supplementary Fig S5A).

Why was no ZC3H20-myc expression detected after induction in bloodstream forms? The results in Fig 6E show that if growing bloodstream forms were treated with tetracycline for 24h, ZC3H20-myc was seen if a protease inhibitor that targets the proteasome was added, but even then the amount was lower than in similarly-treated procyclic forms. ZC3H20-myc is therefore unstable in bloodstream forms, probably being a substrate of the proteasome. In contrast, a preliminary result suggested that ZC3H20-myc has a half-life of at least 6h in procyclic forms (Supplementary Fig S5B).

### Excess ZC3H20 is toxic for procyclic forms

We next asked why cells containing only inducible ZC3H20-myc were unable to grow as procyclic forms. To investigate this, we attempted differentiation in 2.5-fold serial dilutions of tetracycline. After several weeks, a single culture that had been maintained with 1 ng/ml tetracycline started to grow as established procyclic forms. This culture could then grow in the absence of tetracycline. Although ZC3H20-myc was not detectable by Western blotting, it is likely that a small amount was still being expressed. When 100 ng/ml tetracycline was added to these cells, cell growth was strongly inhibited, recovering only when cells lacking detectable expression were selected (Fig 7A). After two days’ expression the cells became swollen (Fig 7A). Importantly, the myc tag was not responsible, because ectopic expression of the unaltered protein had the same effect (Fig 7B). Many of the cells had an enlarged vacuole-like structure between the kinetoplast and the nucleus (Fig 7B, supplementary Fig S6B), which is probably the flagellar pocket. This is reminiscent of the “big-eye” phenotype seen after RNAi targeting clathrin in bloodstream forms (Allen *et al.*, 2003). Incubation with 10ng/ml tetracycline also did not allow growth (Supplementary Fig S5C).

**Fig 7.**
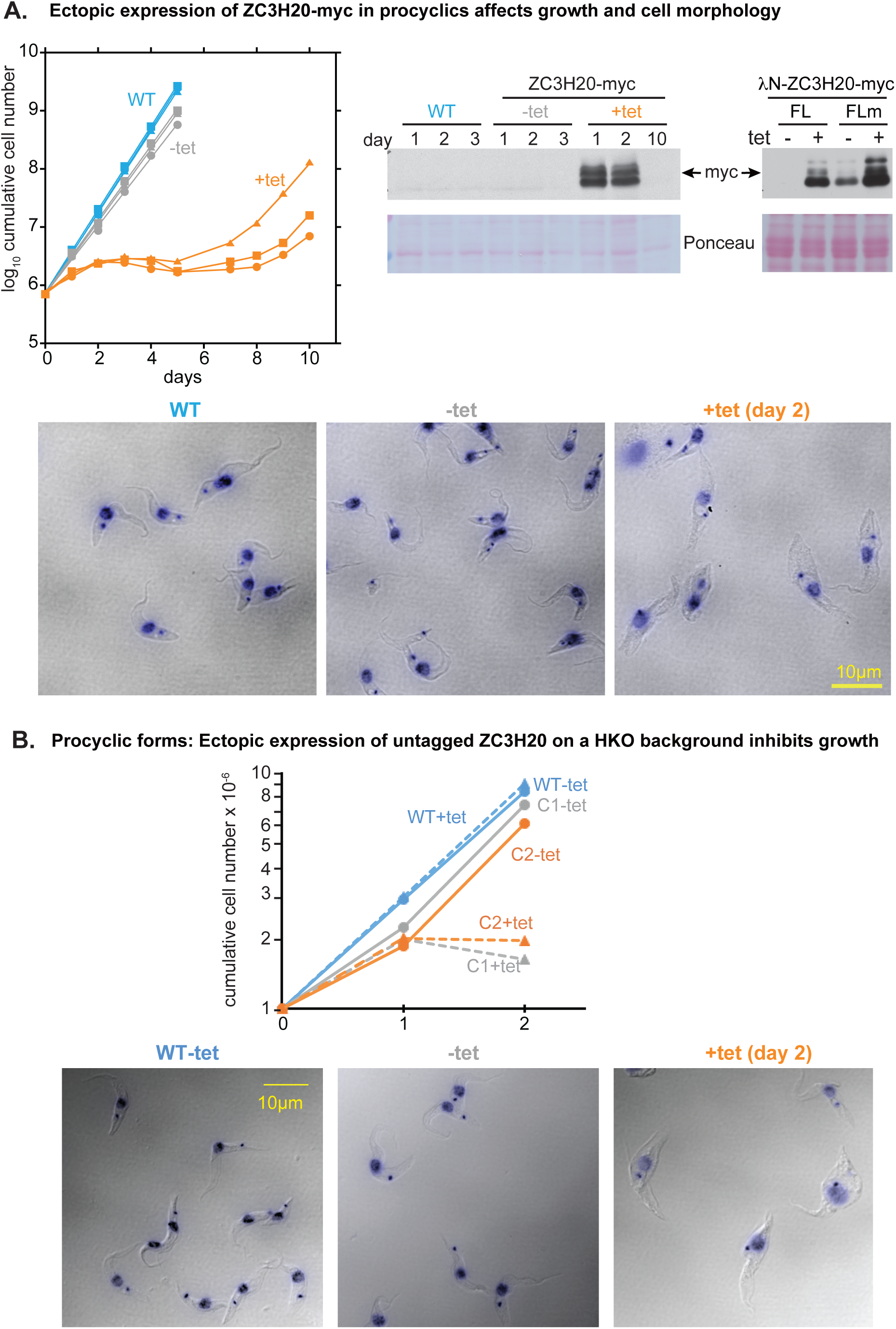
Over-expression of ZC3H20 in procyclic forms is deleterious. A. EATRO 1125 procyclic form cells with inducible ZC3H20-myc and no wild-type genes were treated with 100ng/mL tetracycline to induce expression. Growth curves (three replicates) are on the left and Western blots for expression of ZC3H20-myc, lambdaN-ZC3H20-myc and lambdaN-ZC3H20-myc with a mutant MKT1-binding motif (Fig 5B) are on the right. The micrographs show typical cell morphologies 2 days after induced expression of ZC3H20-myc. Results for lambdaN-ZC3H20-myc were similar whereas the mutant version had no effect. Differential interference contrast images are overlaid DNA stain (DAPI, dark blue). B. EATRO 1125 procyclic form cells with extra inducible ZC3H20 (Supplementary Fig S3 K) were treated with tetracycline to induce the expression. Growth curves for two clones with inducible expression (C1 and C2) are shown above and morphologies of clone 2 cells, on day 2, are below.

To find out whether toxicity of over-expressed ZC3H20 was linked to its ability to promote expression of bound mRNAs, we compared lambdaN-ZC3H20-myc with the HYNPY->HYNAA mutant. LambdaN-ZC3H20-myc was just as toxic as the untagged or myc-tagged version (not shown). The mutant protein was better expressed than the wild-type version (Fig 7A), but had no effect on cell morphology (not shown). This result suggests that toxicity of excess ZC3H20 depends on its ability to activate gene expression, probably via interaction with MKT1.

### ZC3H21 during differentiation and in procyclic forms

The similarities between ZC3H20 and ZC3H21 suggested that they might have shared properties. We therefore wondered whether the absence of ZC3H20 during differentiation might be compensated by premature expression of ZC3H21. We therefore attempted to complement the *ZC3H20* HKO with inducible ZC3H21-myc (Supplementary Fig S3, L). The cells completely failed to differentiate and eventually died (Supplementary Fig S6A). Ectopic expression of ZC3H21 in procyclic forms that retained their *ZC3H21* genes (Supplementary Fig S3, M) caused only a mild growth defect (Supplementary Fig S6B, C) but expression was not maintained, suggesting selection against cells that expressed ZC3H21-myc (Supplementary Fig S6D). Thus ectopic expression of ZC3H21-myc in procyclic forms is probably also deleterious.

### Roles of ZC3H20 and ZC3H21 in procyclic forms

To find out whether either protein is required for growth of procyclic forms, we first depleted each by RNA interference, using cells expressing TAP-tagged proteins. Reduced ZC3H20 in Lister 427 procyclic forms resulted in a very slight growth defect (Fig 2C, Fig 8A) while loss of ZC3H21 had no obvious effect (Fig 8B, Supplementary Fig 3O). We were however unable to generate procyclic forms that completely lacked either ZC3H20 or ZC3H21, despite several attempts, suggesting that such cells are either severely defective or non-viable. This suggests that the two proteins have important, and perhaps essential, independent functions. Loss of ZC3H21 did not result in an increase in TAP-ZC3H20 (Fig 8B).

**Fig. 8.**
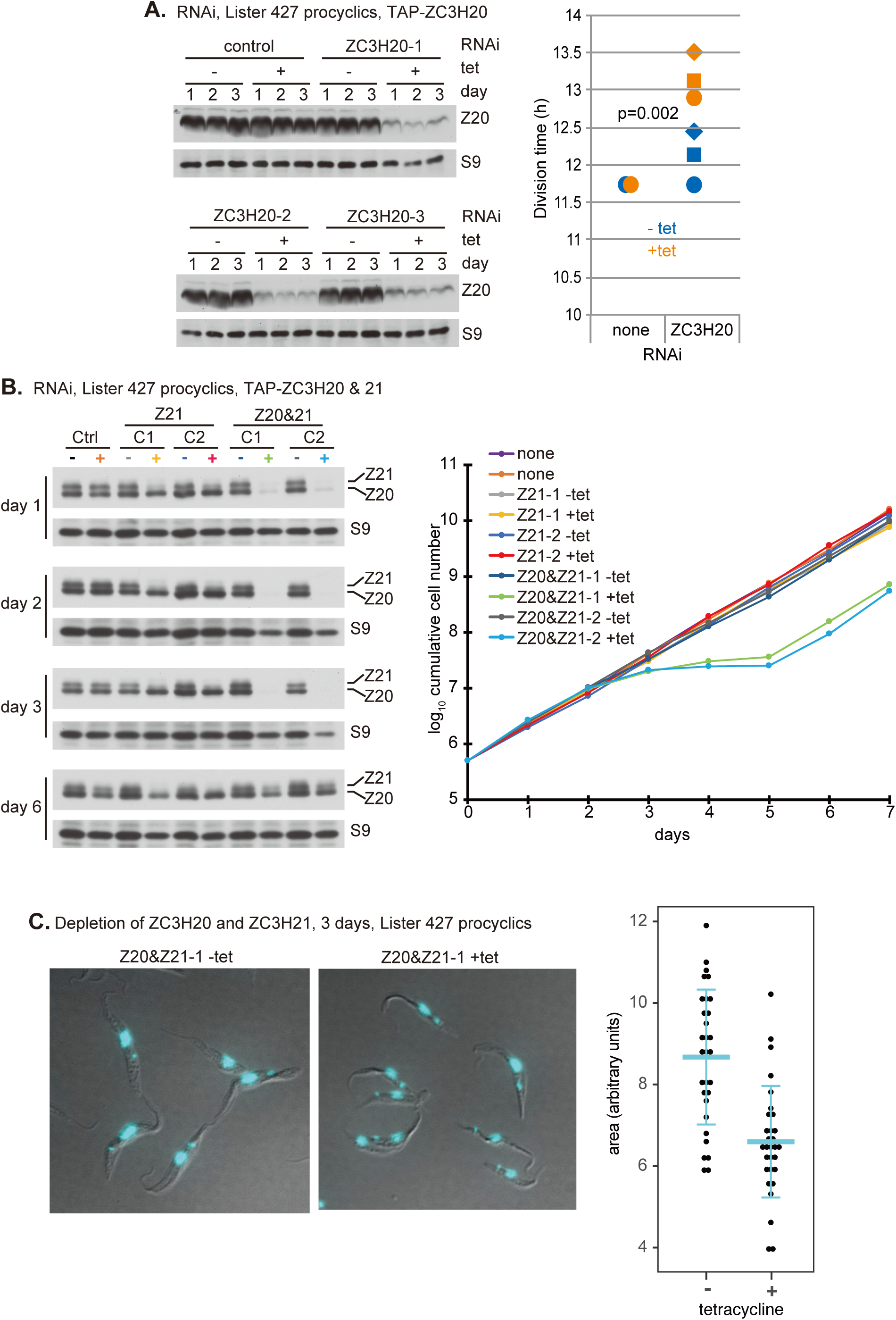
Effects of depleting ZC3H20 or ZC3H20 and ZC3H21 in procyclic forms. A. RNAi targeting *ZC3H20* causes a very slight growth defect. The experiment was done with Lister427 trypanosomes with one *TAP-ZC3H20* gene, one wild-type *ZC3H20* gene, and tetracycline-inducible RNAi (Supplementary Fig S3, panel N). Results are for three independent clones. Expression of TAP-ZC3H20 is shown on the left, and division times are on the right. The p-value is for the difference between +tet and −tet. B. RNAi targeting *ZC3H20* and *ZC3H21*causes a marked growth defect. These cells expressed TAP-ZC3H20 and TAP-ZC3H21, with RNAi targeting a region shared by both genes (Supplementary Fig S3, panel P). Two different RNAi clones were used. The western blots (left) show depletion of both proteins on days 1-3 and recovery of expression on day 6. Growth curves are on the right. C. Left - images of procyclic forms with RNAi targeting *ZC3H20* and *ZC3H21* either without tetracycline, or after 3 days with tetracycline. These are differential interference contrast micrographs with DAPI DNA stain in cyan. The dot plot on the right shows measured cell areas from these and similar images, with mean and standard deviation in cyan.

Simultaneous depletion of both proteins (Supplementary Fig 3P) severely inhibited growth (Fig 8B). After 2 days of RNAi the cells looked relatively normal but after 3 days, they were clearly smaller than non-depleted cells (Fig 8C). We did not see any morphological changes suggesting development of epimastigotes or bloodstream forms. Nevertheless, we tested this possibility by generating EATRO1125 bloodstream forms with the double RNAi, differentiating them to procyclic forms. After 2 weeks, when procyclic-form growth was well established, RNAi was induced. All of the cells died, and could not be rescued by transfer to (tetracycline-containing) bloodstream-form medium either one of two days after RNAi induction (not shown).

### RNAs bound by ZC3H20 and ZC3H21

To find RNA targets of ZC3H20 and ZC3H21, we used cells expressing only TAP-tagged versions (Fig 2B, Supplementary Fig S3I). After UV-cross-linking to stabilise protein-RNA interactions, we purified the proteins on an IgG column, released them with Tobacco Etch Virus protease, and sequenced the associated RNA from duplicate unbound (flow-through) and bound (eluate) fractions (Mugo & Clayton, 2017) (Supplementary Tables S1 and S2). Principal component analysis (Supplementary Fig S7A) indicated that the results were reproducible. Correspondingly, correlations between replicates were reliably very high (R^2^>0.96) (Supplementary Fig S7 B-E) while bound fractions correlated less well with the unbound fractions (Supplementary Fig S7 F-I). In each case, mRNA encoding the target protein was strongly enriched, indicating successful selection of the cognate polysomal mRNA via the N-terminally tagged nascent polypeptide (Supplementary Fig S8A, B). ZC3H20 showed no bias towards longer or shorter mRNAs, whereas ZC3H21 tended not to bind longer mRNAs (Supplementary Fig S8 C, D). The latter effect is only partially due to a lack of binding to mRNAs encoding large cytoskeletal proteins (Supplementary Fig S7 H, I; Supplementary Fig S8D, “cytoskeleton”). When we compared the enriched RNAs, there was moderate correlation between the two datasets (Fig 9A). We defined mRNAs as “bound” if they were reproducibly at least 2-fold enriched in the bound fractions relative to the unbound fractions. 288 mRNAs were reproducibly enriched in the ZC3H20-bound fractions (Supplementary Table S1, sheet 1), and 238 were ZC3H21-bound (Supplementary Table S2, sheet 1). Of these, 101 were shared (Fig 9A). This confirmed our prediction that the two proteins might have overlapping, but not identical, specificities. Both ZC3H20 and ZC3H21 showed a clear preference for mRNAs encoding possible membrane proteins and mitochondrial proteins (Fig 9C and Supplementary Fig S8 C, D). Quantitative comparisons for all functional classes (Supplementary Fig S9) also showed biases towards mRNAs encoding proteins of the plasma membrane and mitochondrial membrane. ZC3H20 showed no binding to mRNAs encoding ribosomal proteins (Supplementary Fig S10).

**Fig. 9.**
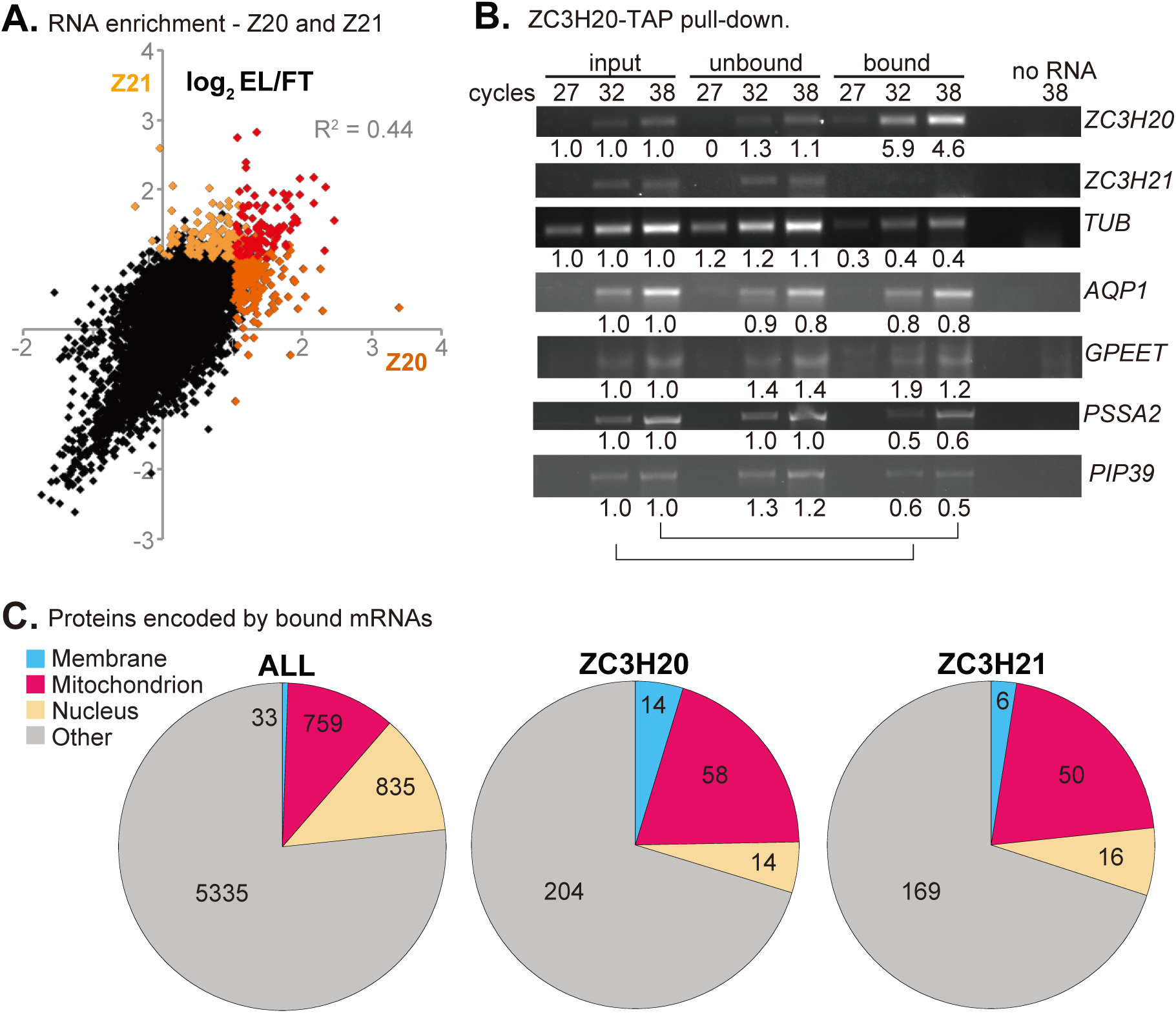
Binding of ZC3H20 and ZC3H21 to mRNAs. A. ZC3H20 and ZC3H21 show overlapping mRNA binding specificities. The average binding ratios (RPM of eluate / unbound fractions) for each mRNA are plotted for ZC3H20 (x-axis) and ZC3H21 (y-axis). A list of unique mRNAs ((Siegel *et al.*, 2010) and see Supplementary Tables) was used for the analysis. Bound mRNAs were defined as those that gave ratios of at least 2-fold in both experiments. mRNAs bound by both proteins are in red, those bound only by ZC3H20 are in dark orange and by ZC3H21 in pale orange. B. Verification of specific mRNA binding by ZC3H20. RNA was pulled down and subjected to RT-PCR. Products were analysed by agarose gel electrophoresis with ethidium bromide staining. Results for various cycle times are shown. Numbers beneath the lanes are signal relative to input with the same number of cycles (as indicated by the lines at the bottom). The data are semi-quantitative and the intensity differences between cycles were lower than expected. As expected, *ZC3H20* mRNA was pulled down, presumably via the nascent polypeptide, while the *ZC3H21* mRNA was not. C. Locations of proteins encoded by mRNAs bound by ZC3H20 and ZC3H21. Locations were assigned using information in TritrypDB, TrypTag (Dean *et al.*, 2016) and various publications. The pie chart on the left shows locations of all proteins encoded by the unique gene list.

Membrane protein mRNAs that were bound to both proteins encoded amastin-like protein, trans sialidases, GPI-anchored membrane proteins including GPEET procyclin, a flagellar membrane protein, and a procyclic surface protein (PSSA-2). ZC3H21 also was found to bind the *EP2* mRNA; binding by ZC3H20 was just below our cut-off. The two mRNAs that Ling et al. (Ling *et al.*, 2011) had previously shown to be bound to ZC3H20, encoding a *trans* sialidase (Tb927.5.440) and the mitochondrial carrier MCP12 (Tb927.10.12840), were bound to both ZC3H21 and ZC3H20 in our analysis. We independently confirmed binding of ZC3H20 to several the mRNAs, encoding Aquaporin 1, GPEET procyclin, and procyclic-specific surface antigen 2, by pull-down followed by semi-quantitative PCR (Fig 9B).

Since several of the identified genes belong to multi-gene families, we decided to distinguish between the members. To do this we set the alignment criteria such that all reads that mapped to the genome more than once were eliminated. Normalisation of these results is problematic because most abundant mRNAs are encoded by at least two genes: repeated sequences constitute 80-95% of reads (Supplementary Tables S1 and S2, sheets 5 and 6, MAPQ42 alignments). For ZC3H21 in particular, normalisation of the MAPQ42 results to reads per million reads yielded results that contradicted those obtained when entire mRNAs were considered (Supplementary Table S2, sheets 5 & 6). Nevertheless, for ZC3H20, the results did suggest selective enrichment of several additional mRNAs, encoding various amino acid transporters, ESAG9 variants, a META domain protein, and PAD2 (Supplementary Table S1, sheet 1). Binding to both *EP* and *GPEET* mRNAs was also revealed (Supplementary Table S1 and S2, sheet 1; Supplementary Figs S10 and S11). Neither protein bound to the mRNA encoding PAD1, and ZC3H21 did not bind trans sialidase mRNAs. Of the six mRNAs that were previously reported as decreased after ZC3H20&21 RNAi (Ling *et al.*, 2011), we identified four as being bound to ZC3H20: META domain, trans sialidase, neuraminidase, and MCP12. To examine developmental regulation of the bound mRNAs, we concentrated on the numbers of ribosome footprints, since these measure both mRNA levels and the densities of bound ribosomes (Antwi *et al.*, 2016). Across the whole dataset, there was no correlation between ZC3H20 or ZC3H21 binding and developmental regulation (Supplementary Fig S8 A, B). Although several highly enriched mRNAs have higher ribosome occupancy counts in procyclic forms, both proteins also appeared to bind unregulated, or even bloodstream-form-specific mRNAs (Supplementary Fig S8 A, B, and Supplementary Tables S1 and S2, sheet 1).

Each zinc finger of ZC3H20 or ZC3H21 is expected to bind to four nucleotides (Hudson *et al.*, 2004). We used DREME to search for motifs that were present in bound mRNAs (Supplementary Tables S1 and S2, sheet 1), and absent in those that were depleted (eluate lower than the unbound fraction) (Supplementary Tables S1 and S2, sheet 3). No convincing motifs were found.

### Effects of ZC3H20 and ZC3H21 depletion on the transcriptome

To find out how loss of ZC3H20 impacted bloodstream-form trypanosome differentiation, we induced RNAi 12h after cis-aconitate addition and temperature reduction (Fig 6A). Cells were harvested 24h later and mRNA subjected to RNASeq (Fig 6A, Supplementary table S3). All differences were relatively minor, and they mostly reflected the fact that cells lacking ZC3H20 were failing to switch towards a procyclic-form expression pattern. *ZC3H20* reads were reduced only 2-fold, suggesting poor RNAi induction. Despite this, the results were informative. The three other mRNAs that were more than 1.5-fold significantly decreased (Supplementary table S3, sheet 1) encode a trans-sialidase, flabarin, and the heme uptake protein LHR1. The amastin, GPEET, PSSA-2 and MCP12 mRNAs were also at least 1.3-fold lower after RNAi than in the control. Of the fifty-one mRNAs with at least 1.3-fold reductions, nineteen had been shown to bind to ZC3H20. In contrast, five ZC3H20-bound mRNAs were that encode proteins of unknown function increased relative to the controls (Supplementary table S3, sheet 1).

The effects of depleting both ZC3H20 and ZC3H21 in procyclic-form Lister 427 cells were stronger. This is essentially a repeat of the experiment done by Ling et al. (Ling *et al.*, 2011), but using the much more sensitive RNASeq technology and a slightly shorter RNAi induction time (36h instead of 48h) to reduce secondary effects. 136 mRNAs were more than 1.5-fold decreased, and 110 more than 1.5-fold increased (Supplementary Table S4). Many of the changes reflected a shift towards bloodstream-form or metacyclic-form metabolism (Fig 10A), and the results also correlated, to some extent, with those seen after ZC3H20 RNAi in differentiating cells (Fig 10B) (Supplementary Tables S3, S4). We confirmed four of the five mRNAs that had been identified by Ling et al. (Ling *et al.*, 2011) as being significantly reduced: trans-sialidases, mitochondrial carrier protein MCP12, Tb927.5.4020, and GRESAG4. The remaining genes, members META gene family, were not in our unique gene list but were also increased in the raw dataset. The ten increased mRNAs from Ling et al showed a similar overlap with our data.

**Fig. 10.**
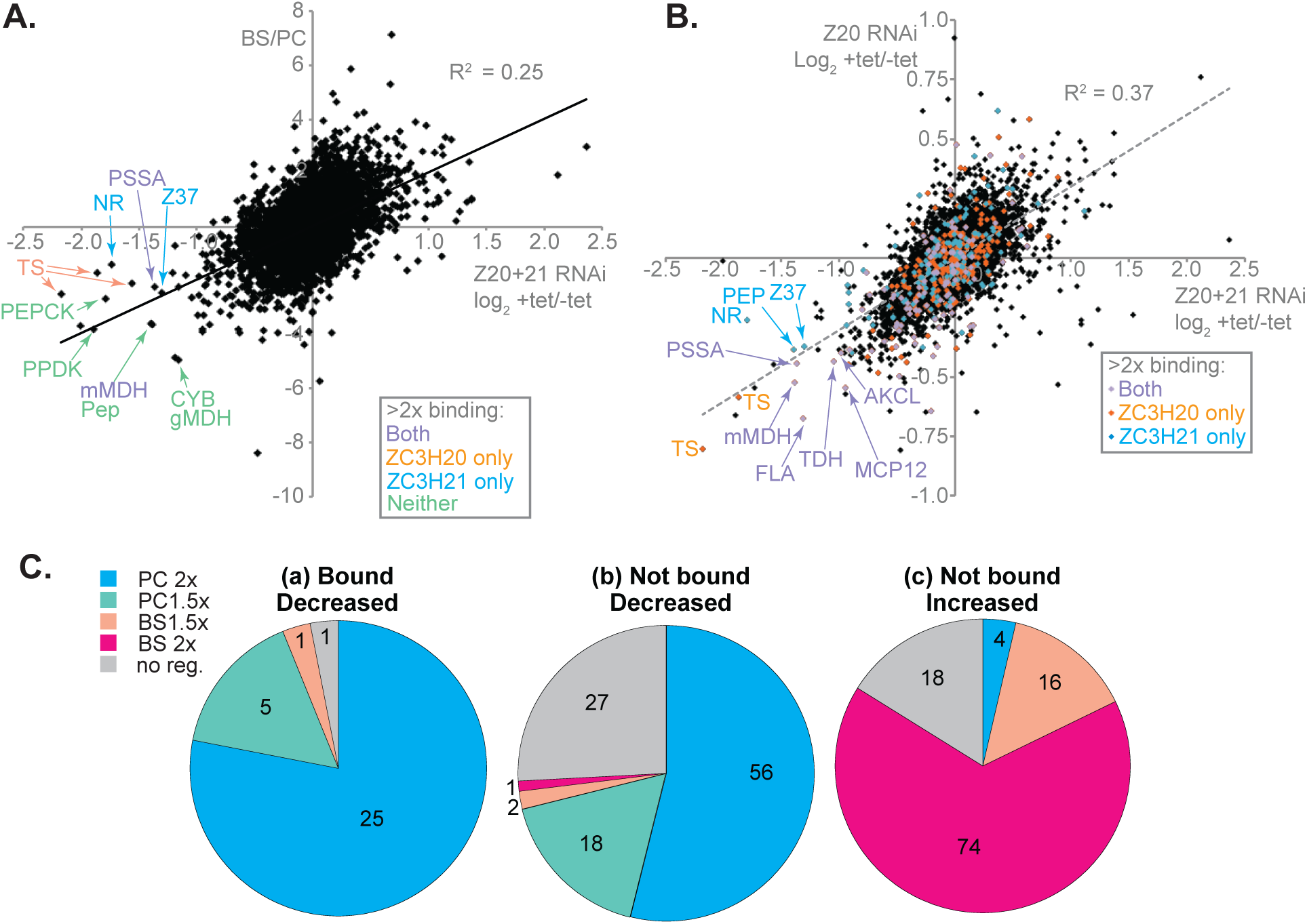
Effects of depleting ZC3H20 only, or ZC3H20 and ZC3H21 simultaneously, on the transcriptome. RNAi against *ZC3H20* was induced in EATRO1125 bloodstream-form trypanosomes incubated at high density (3×10^6^/ml) for 10h, then incubated for 24h at 27*C with tetracycline and cis-aconitate prior to RNA preparation. RNAi to deplete both proteins in Lister 427 procyclic forms was induced for 36h. A. Correlation between the effects of *ZC3H20* and *ZC3H21* RNAi in procyclic forms, and developmental regulation of mRNA (RNA amounts in bloodstream forms divided by the amount in procyclic forms). Some of the strongly decreased mRNAs are indicated, and colour-coded according to their binding to ZC3H20 and/or ZC3H21 (key on Fig). Labelled spots for chosen genes are as follows: AKCL - 2-amino-3-ketobutyrate coenzyme A ligase; FLA - flabarin; MDH - Malate dehydrogenase (m-mitochondrial, g-glycosomal); MCP12: mitochondrial carrier protein 12; NR - Nitrate reductase; Z37 - ZC3H37; TDH - L-threonine 3-dehydrogenase; Pep - peptidase; PEPCK: phosphoenol pyruvate carboxykinase; PPDK: pyruvate phosphate dikinase; PSSA - Procyclin specific surface antigen; TS - trans sialidase; Z37: ZC3H37. All of these mRNAs were reduced after *ZC3H20* RNAi. B. Correlation between the effects of *ZC3H20* and *ZC3H21* RNAi in procyclic forms (x-axis) and the effects of *ZC3H20* RNAi in differentiating cells (y-axis). Coloured dots indicate RNAs that were bound by either protein (key on Fig). Abbreviations for labelled spots are as in (A). C. Developmental regulation of mRNAs that showed at least a 1.5-fold change after RNAi targeting both ZC3H20 and ZC3H21 in procyclic forms. (a) Results are shown for 32 bound RNAs that decreased at least 1.5 x after RNAi. These genes are a subset of those in Supplementary Table S5, sheet 1. Of the 436 mRNAs that were bound to either or both proteins, 8 also increased >1.5× after RNAi. Of these, one is 1.5× more abundant in bloodstream forms than in procyclics. (b) and (c) include results for all mRNAs that were not enriched at least 2-fold in both RNA-pull-down preparations for either ZC3H20 and ZC3H21. (A few of these may in fact be bound, but did not satisfy the cut-off criterion). These genes are listed in Supplementary Table S5, sheet 2.

**Fig. 11.**
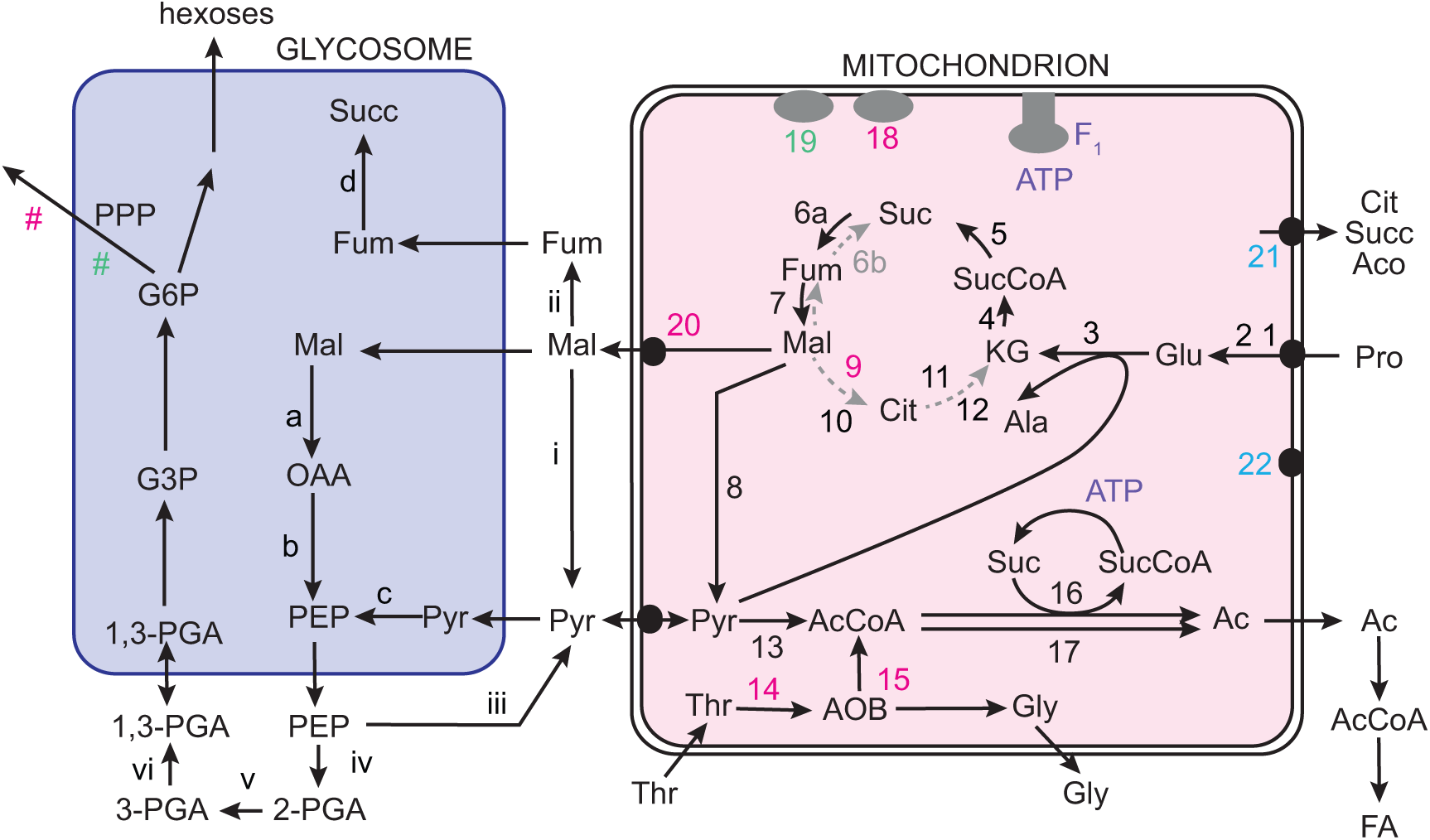
Effects of depleting ZC3H20 and ZC3H21 on energy metabolism. Information for this Fig is from (Allmann *et al.*, 2013, Coustou *et al.*, 2008, Wargnies *et al.*, 2018, Millerioux *et al.*, 2013). The black arrows indicate metabolism in cells grown in the proline-rich, glucose-poor medium used in our study. Dotted lines are steps that are possible but may not be active under these conditions. The presence of a complete citric acid cycle has not been demonstrated under any conditions. Numbers in magenta are enzymes for which the mRNAs are bound by both ZC3H20 and ZC3H21, and which also decrease after the double RNAi. Those in cyan are bound by ZC3H20 and decreased after RNAi; green indicates binding and regulation by ZC3H21. Succinate dehydrogenase (6a) is essential in the absence of glucose (Coustou *et al.*, 2008) Abbreviations are: PPP-pentose phosphate pathway; G6P - glucose-6-phosphate; G3P - glyceraldehyde-3-phosphate; PGA - phosphoglycerate; Suc - succinate; Fum - fumarate; Mal - malate; OAA - oxaloacetate; PEP - phosphoenol pyruvate; Pyr - pyruvate; KG - alpha-ketoglutarate; Cit - citrate; Gly - glycine; Pro - proline; Aco - aconitate; AOB - amino oxobutyrate; Thr - threonine; Gly - glycine; Ac - acetate; FA - fatty acids; AcCoA - acetyl CoA. Enzymes are: (1) proline dehydrogenase; (2) Δ1-pyrroline-5-carboxylate dehydrogenase ; (3) L-alanine aminotransferase; (4) a-ketoglutarate dehydrogenase; (5) succinyl CoA synthase; (6a) succinate dehydrogenase (complex II); (6b) m fumarate reductase ; (7) mitochondrial Fumarase; (8) mitochondrial malic enzyme; (9) m malate dehydrogenase; (10) citrate synthase; (11) aconitase; (12) isocitrate dehydrogenase (NADP); (13) pyruvate dehydrogenase complex; (14) L-threonine 3-dehydrogenase; (15) 2-amino-3-ketobutyrate CoA ligase; (16) acetate:succinate CoA transferase; (17) acetyl coA thioesterase; (18) cytochome c oxidase complex; (19) cytochrome b5 reductase; (20) MCP12; (21) tricarboxylate carrier?; (22) POMP1; (i) cytosolic malic enzyme; (ii) cytosolic fumarase; (iii pyruvate kinase; (iv) enolase; (v) phosphoglycerate mutase; (vi) cytosolic phosphoglycerate kinase; (a) glycosomal malate dehydrogenase; (b) phosphoenol pyruvate carboxykinase; (c) pyruvate phosphate dikase; (d} fumarase/ fumarate hydratase; # pentose phosphate pathway (PPP) enzymes.

To find direct effects of ZC3H20 and ZC3H21 depletion, we divided the affected mRNAs according to whether or not they were bound by ZC3H20 or ZC3H21, and focused on mRNAs that decreased after RNAi (Supplementary Table S5, sheet 1 and Table 1). The vast majority of bound mRNAs were not affected by RNAi (Fig 10B). This could be because the depletion time was too short, depletion was insufficient, or other bound RNA-binding proteins override the effects of loss of ZC3H20 and ZC3H21. Nearly all of the bound mRNAs that decreased after RNAi are procyclic-specific (Fig 10 C (a)). They are enriched for location to the plasma membrane (P = 7 × 10^-6^, Fisher test adjusted for multiple testing) or the mitochondrion (P = .018). Analysis of the 3’-UTRs from these mRNAs, in comparison with a set bound by neither protein, revealed a potential enriched motif, (G/U)UUA(U/G)CG. However this motif was only found in about half of the bound and regulated mRNAs so its significance is unclear.

**Table 1.** Mitochondrial and membrane protein mRNAs that are bound and regulated by ZC3H20 and ZC3H21. Transcripts were classified as bound only if they were at least two-fold enriched in both replicates. For regulation the threshold was much less stringent (at least a 1.3× decrease, Padj <0.05). “BS/PC ribosome profiling” shows the ratios of normalised total ribosomal reads (Antwi *et al.*, 2016) and “BS/PC RNA” is regulation at the RNA level (Fadda *et al.*, 2014). Nearly all affected mRNAs shown are preferentially expressed in procyclic forms. Additional mRNAs are bound and regulated, but do not encode membrane or mitochondrial proteins.

| Gene ID -<br>Tb927 | Annotation | Fold<br>decr<br>ease<br>,<br>Z20_<br>21<br>RNAi | Fold<br>decr<br>ease<br>, Z20<br>RNAi | Z20<br>boun<br>d/<br>unbo<br>ud | Z21<br>boun<br>d/<br>unbo<br>ud | BS/P<br>C<br>ribos<br>ome<br>profi<br>ling | BS/<br>PC<br>RNA |
| --- | --- | --- | --- | --- | --- | --- | --- |
| **Membrane protein mRNAs bound by ZC3H20 and 21** |  |  |  |  |  |  |  |
| 4.3500 | Amastin surface glycoprotein | 1.64 | 1.32 | 3.64 | 2.84 | 0.11 | 0.35 |
| 6.510 | GPEET2 procyclin | 1.39 | 1.41 | 4.51 | 4.52 | 0.00 | 0.01 |
| 10.11220 | procyclic form surface glycoprotein | 2.56 | 1.35 | 3.34 | 3.84 | 0.09 | 0.28 |
| 11.2410 | Flabarin flagellar membrane protein | 2.5 | 1.59 | 3.14 | 2.72 | 0.01 | 0.07 |
| 7.4270 | Unknown function | 1.39 | 1.23 | 2.94 | 2.75 | 0.23 | 0.41 |
| **Membrane protein mRNAs bound by ZC3H20** |  |  |  |  |  |  |  |
| 8.1620 | MSP-B | 1.64 | 1.23 | 2.43 | 2.29 | 0.05 | 0.26 |
| 5.440 | trans sialidase | 4.55 | 1.75 | 2.86 | 1.18 | 0.00 | 0.04 |
| 8.7340 | trans-sialidase,neuraminidase | 3.7 | 1.49 | 2.13 | 1.34 | 0.02 | 0.13 |
| 6.1520 | Aquaglyceroporin 1 | 1.37 | 1.33 | 3.07 | 1.42 | 0.16 | 0.51 |
| 11.1830 | flagellar membrane protein | 1.33 | 1.35 | 3.17 | 1.33 | 0.17 | 0.48 |
| **Mitochondrial protein mRNAs bound by ZC3H20 and 21** |  |  |  |  |  |  |  |
| 10.510 | ATOM19 | 1.32 | 1.16 | 2.18 | 2.21 | 0.34 | 0.44 |
| 9.12500 | POMP7 | 1.45 | 1.12 | 3.33 | 2.75 | 0.22 | 0.62 |
| 10.12840 | mitochondrial carrier protein MCP12 | 1.92 | 1.47 | 2.28 | 2.1 | 0.09 | 0.22 |
| 10.2560 | mitochondrial malate dehydrogenase | 2.63 | 1.43 | 2.55 | 7.1 | 0.05 | 0.21 |
| 6.2790 | L-threonine 3-dehydrogenase | 2.04 | 1.35 | 2.34 | 2.19 | 0.30 | 0.62 |
| 8.6060 | 2-amino-3-ketobutyrate coenzyme A ligase | 1.96 | 1.32 | 2.32 | 4.07 | 0.36 | 0.70 |
| 10.3120 | cytochrome c oxidase assembly protein | 1.37 | 1.22 | 2.12 | 2.35 | 0.29 | 0.71 |
| 10.13600 | component of cytochrome oxidase complex? | 1.3 | 1.18 | 2.98 | 2.34 | 0.34 | 0.86 |
| 1.1000 | Phosphoprotein dr6 | 1.45 | 1.11 | 4.48 | 3.51 | 0.63 | 0.76 |
| 3.3330 | HSP20, heat shock protein 20 | 1.54 | 1.28 | 3.33 | 2.81 | 0.40 | 0.63 |
| 9.6620 | Unknown function | 1.33 | 1.14 | 2.2 | 3.1 | 0.10 | 0.49 |
| 8.2470 | Unknown function | 1.69 | 1.04 | 5.05 | 4.1 | 0.27 | 0.34 |
| **Mitochondrial proteins mRNAs bound by ZC3H20 only** |  |  |  |  |  |  |  |
| 9.2320 | POMP1 - S-Adenosyl methionine methyl transferase? | 1.59 | 1.39 | 2.96 | 1.28 | 0.11 | 0.54 |
| 9.4310 | Tricarboxylate carrier? | 1.89 | 1.37 | 2.3 | 2.01 | 0.11 | 0.33 |
| 11.3750 | NADH-cytochrome b5 reductase | 1.35 | 1.09 | 2.29 | 1.9 | 0.06 | 0.41 |
| 11.7900 | mitochondrial RNA binding protein 16 (RBP16) | 1.47 | 1.27 | 2.92 | 2.16 | 0.34 | 0.38 |
| 6.2390 | Unknown function | 1.45 | 1.09 | 2.12 | 1.29 | 0.54 | 0.77 |
| **Mitochondrial protein mRNAs bound by ZC3H21 only** |  |  |  |  |  |  |  |
| 5.3040 | Cytochrome c oxidase complex, MIX protein | 1.41 | 1.22 | 1.93 | 2.53 | 0.19 | 0.54 |
| 7.2700 | NADH-cytochrome b5 reductase | 1.89 | 1.09 | 1.6 | 2.27 | 0.14 | 0.36 |
| 11.1310 | NADH-cytochrome b5 reductase | 1.49 | 1.09 | 1.94 | 2.58 | 0.61 | 0.80 |
| 4.1670 | POMP29, cytochrome b5-like haem-binding | 1.89 | 1.15 | 1.41 | 2.89 | 0.15 | 0.46 |
| 11.16480 | enoyl-CoA hydratase/isomerase family protein | 1.92 | 1.22 | 1.64 | 3.48 | 0.11 | 0.29 |
| 11.13510 | Unknown function | 1.45 | 1.19 | 1.9 | 2.09 | 0.20 | 0.54 |

The mRNAs that were not bound by either ZC3H20 or ZC3H21, but nevertheless showed a significant decreases after RNAi, were predominantly procyclic-specific (Fig 10C(b) and Supplementary Table S5, sheet 2), while those that increased after RNAi were mainly bloodstream-form specific (Fig 10C(c)). Thus loss of ZC3H20 and ZC3H21 target mRNAs precipitated a change in the transcriptome towards the bloodstream-form pattern. Expression of RBP6 induces differentiation of procyclic forms to epimastigotes, and thence to rodent-infective metacyclic trypomastigotes; and expression of RBP10 switches procyclic forms to metacyclic-like forms that can proliferate as bloodstream forms. The mRNAs encoding both of these proteins were increased after *ZC3H20+21* RNAi. There were also increases in mRNAs encoding three other bloodstream-form specific RNA-binding proteins: RBP26, ZC3H32, and HNRNPH/F, and in mRNAs encoding the epimastigote coat protein BARP (Urwyler *et al.*, 2007) and HAP2, which is implicated gamete fusion (Peacock *et al.*, 2011). The effects of *ZC3H20+21* RNAi were however only very partially similar to those seen after ectopic expression of RBP10 (Supplementary Fig S13) and as noted above, the cells could not grow as bloodstream forms. Loss of ZC3H20 and 21 target mRNAs may therefore be too deleterious to allow further differentiation.

## Discussion

The results we have described indicate that ZC3H20 and ZC3H21 are mRNA-binding proteins with overlapping, but non-redundant functions. ZC3H20 is expressed at a low level in bloodstream forms, reaches a maximum in stumpy forms, then persists in procyclic forms. Only a minor part of this regulation is at the mRNA level, perhaps mediated by RBP10 (this paper): *ZC3H20* ribosome footprints are only 2-fold less abundant in bloodstream forms than in procyclic forms (Antwi *et al.*, 2016). The remaining regulation is accounted for by protein degradation in bloodstream forms. ZC3H21 shows slightly stronger (3-fold) regulation of ribosome footprints, and appears only after differentiation has been initiated. Phosphorylation of both proteins has been detected (Urbaniak *et al.*, 2013), but whether this has any role in their function or stability is not known. The difference in behaviour of the *ZC3H20* and *ZC3H21* mRNAs might partially be explained by the fact that the *ZC3H21* 3’-UTR has five RBP10 binding sites, while that of *ZC3H20* has only two. As RBP10 levels decrease during differentiation, binding to *ZC3H20* will be lost first.

The products of mRNAs that are both bound and stabilised by ZC3H20 and ZC3H21 were strongly enriched in proteins of the plasma membrane and mitochondrion, and all of those mRNAs are more abundant in procyclic than in bloodstream forms (Supplementary Table S5, sheets 1 and 2). Stabilisation of membrane protein mRNAs may explain why the cells become smaller after *ZC3H20* and *ZC3H21* RNAi (Fig 8C), and swollen upon ZC3H20 over-expression (Fig 7 and Supplementary Fig S6). Although procyclic forms can grow and divide without EP and GPEET procyclins (Vassella *et al.*, 2003), they may not be able to do so if they also lack trans sialidases, PSSA-2, and amastin. Conversely, ZC3H20 over-expression might result in an overload of the secretory pathway and of protein transfer from the flagellar pocket to the outer plasma membrane, resulting in swelling of the flagellar pocket; trans sialidases, which are regulated by ZC3H20 but not ZC3H21, seem the most likely culprits, but flabarin and another flagellar membrane protein (Tb927.11.1830), might also be implicated.

Metabolic effects might also contribute to the small size of ZC3H20 and ZC3H21-depleted cells. The RNAi caused decreases in mRNAs encoding two enzymes required for threonine conversion to acetyl coA (Fig 10) - a major source of acetate for lipid biosynthesis (Millerioux *et al.*, 2013). Moreover, mitochondrial energy metabolism should be impacted, because of decreases in di- or tri-carboxylate carriers, the cytochrome oxidase and cytochrome b5 reductase complexes, and mitochondrial malate dehydrogenase (Fig 10). As an indirect measure of ATP status, we measured the extent to which the cells could swim above the bottom of the flasks, but observed no effect of RNAi (not shown), so alternative pathways may compensate; or perhaps ATP is available for movement because it is not being used for cell growth. Finally, ZC3H20 and ZC3H21 target mRNAs encoding five proteins that are implicated in pre-rRNA processing, and whose loss could result in slower growth.

ZC3H20 is required for stumpy form generation. There are several possible reasons for this, which might act alone or in combination. One mRNA that was mildly decreased in cells with ZC3H20 RNAi after 24h cis-aconitate encodes PIP39, a protein phosphatase. *PIP39* mRNA is not developmentally regulated but the protein has increased abundance in stumpy forms and is required for their formation (Szoor *et al.*, 2010). *PIP39* mRNA had moderate binding to ZC3H20 (just below our chosen threshold) so might require ZC3H20 for either stabilization, or increased translation. Loss of stumpy formation might also be caused by a failure to up-regulate expression of mitochondrial proteins. However, stumpy forms can metabolise glucose (Vickerman, 1965) and neither ATP generation by the F_1_ ATPase nor the membrane potential are required (Dewar *et al.*, 2018), which suggests that they do not need to import newly-synthesized proteins. Another possibility is that some of the plasma membrane proteins are needed. Several of them are increased in stumpy forms at both RNA (Silvester *et al.*, 2018) and protein (Dejung *et al.*, 2016) levels, including trans sialidases, amastin-like protein, flabarin, and PSSA-2. Possible roles for proteins that currently have no known function (Tb927.11.16300, Tb927.11.3830) can of course never be excluded, and in some cases, loss of ZC3H20 may affect translation without an immediate impact on mRNA abundance. For example, the NrkA protein kinase encoded by Tb927.4.5390 (an mRNA that is also bound by RBP10 (Mugo & Clayton, 2017) and is increased in stumpy forms (Silvester *et al.*, 2018)) clearly warrants further investigation. Its depletion inhibited differentiation in a high-throughput screen (Alsford *et al.*, 2005).

The binding specificities of ZC3H20 and ZC3H21 were not directly investigated here and the only possible consensus sequence, (G/U)UUA(U/G)CG, was missing in half of the bound and regulated transcripts. Two regions of the EP procyclin 3’-UTR are implicated in mRNA stability and translation in procyclic forms: the first 40 nt, and a 16mer stem-loop towards the 3’-end (Hehl *et al.*, 1994). The 16mer loop sequence is CCCUGUAGAUUU, which does not match the putative consensus. The procyclin mRNAs are destabilised by RBP10, which binds to a different part of the 3’-UTR. Thus as RBP10 decreases, and ZC3H20 increases, the balance shifts from degradation towards translation and stability. The later appearance of ZC3H21 may further promote procyclin expression, particularly of EP2 and EP3. The procyclin mRNAs are not the only ones to be subject to this dual regulation: RBP10 binds to 4% of all mRNAs, but to14% of all mRNAs that are bound by ZC3H20 and/or ZC3H21 (Supplementary Table S5, sheet 1).

In addition to changes in direct target mRNAs, depletion of ZC3H20/21 resulted in changes in expression of numerous mRNAs that were not bound by either protein. Many of these changes reflected a shift towards a more bloodstream-form transcript pattern. We suggest that they are likely to be secondary effects. One possibility is a response to stress, although there was no correlation with the effects of heat shock (Minia *et al.*, 2016) or starvation (Fritz *et al.*, 2015). Altered metabolite levels could also be responsible (Vassella *et al.*, 2004, Vassella *et al.*, 2000, Bringaud *et al.*, 2006, Qiu *et al.*, 2018, Fernandez-Moya *et al.*, 2014); and it is conceivable that one or more of the directly regulated surface proteins acts as a receptor for a signal that maintains procyclic identity.

We conclude that during differentiation of bloodstream forms to procyclic forms, loss of RBP10, and of a specific degradation mechanism, causes an increase in ZC3H20, enabling trypanosomes to prepare for growth as procyclics by stabilising mRNAs that encode mitochondrial enzymes and procyclic surface proteins. e suggest that, in mRNAs that bear binding sites for both proteins, the appearance of ZC3H20 within the mRNP diminishes the influence of RBP10, preventing mRNA destruction. During full differentiation to procyclics and following complete loss of RBP10, ZC3H21 appears, reinforces the effects of ZC3H20 and stabilises additional mRNAs. Our results thus not only illuminate mechanisms of mRNA control during trypanosome differentiation, but also provide an excellent general example of how mRNAs with more than one specific protein binding site can be subject to stringent control through changing levels of cognate binding proteins with competing activities.

## Experimental procedures

### Trypanosome culture

Bloodstream-form *T. brucei brucei* 427 Lister strain were cultured in HMI-9 plus 10% foetal calf serum, Procyclic form *T. b. brucei* were cultured in MEM-Pros plus 10% foetal calf serum at 27°C. Tetracycline-regulated transcription was induced with of 0.1µg/ml tetracycline.

Stable cell lines were generated as described in (Benz *et al.*, 2011). All plasmid constructs and oligonucleotides are listed in Supplementary Table S6.

For differentiation, the pleomorphic EATRO 1125 cell lines were growing in medium containing 1.1%methyl cellulose to around 3×10^6^ cells/mL, then maintained at this density for 10-14h. PAD1 expression was checked by Western blot. Differentiation was triggered by adding 6mM cis-aconitate and lowering the temperature to 27°C. After 24h the medium was changed to MEM-Pros.

### Protein and Northern blot analyses

For protein analysis, 2-3×10^6^ cells were collected per each sample, resuspended in Laemmli buffer heated and subjected to SDS-PAGE gel electrophoresis. All assays of macromolecular biosynthesis and RNA processing were done at densities of less than 2 × 10^6^/ml. Antibodies used for Western blots were: HRP conjugated anti-peroxidase (1:5000, Sigma); rat anti RBP10 (1:2000) ; mouse anti-myc (1:1000 Santa Cruz); mouse anti EP(1:1000, Cedar Lane); rat anti S9 (1:2000); rabbit anti-PAD1(1:1000) (kind gift from Keith Matthews). HRP conjugated second antibodies (GE Healthcare) were used at a concentration of 1:5000. Detection was done using Western Lightning-Plus (Perkin-Elmer) followed by exposure using X-ray films.

Total RNA was extracted from roughly 5×10^7^ cells using peqGold TriFast (peqLab) following the manufacturer’s instructions. The RNA was separated on formaldehyde gels and then blotted on Nytran membranes (GE Healthcare). Following crosslinking and methylene blue staining (SERVA). For mRNA detection, the membranes were incubated with [α-^32^P]dCTP radioactively labelled DNA probes (Prime-IT RmT Random Primer Labelling Kit, Stratagene) overnight at 65°C. After washing the blots, they were exposed to autoradiography films and detection was performed with FLA-7000 (GE Healthcare). The images were processed and quantified with Fiji (ImageJ) and Adobe Photoshop.

All tethering assays were done as described in (Mugo & Clayton, 2017).

### RNA pull-down and sequencing

mRNAs bound to ZC3H20 and ZC3H21 were identified as described previously (Droll *et al.*, 2013, Mugo & Clayton, 2017). Two independent purifications were done for each protein, using EATRO1125 procyclic forms expressing only TAP-tagged versions of the relevant protein. In each replicate, 1-2×10^9^ cells were centrifuged down and resuspended in 25mL 4°C cold PBS, then put in a 145mm-diameterpetri dish on ice. The RNA-protein complexes were crosslinked by twice irradiating the cells with UV at 240 mJ/cm^2^. All procedures were performed at 4°C or on ice unless otherwise stated. The cells were then centrifuged washed once with ice-cold PBS. After removing the supernatant the pellets were snap frozen in liquid nitrogen. Cell pellets can be stored in -70°C for several days before the RNA pull-down experiment. For the RNA pull-down, cells were broken in 1mL lysis buffer (20mM Tris-HCL PH7.5, 5mM MgCl_2_, 0.1% IGEPAL GA-630, 1mM DTT, 100U RNase inhibitor, proteinase inhibitor cocktail) by passaging through syringe needled (20 times 21G1×1/2 then 20 times 27Gx 3/4). Cell debris was pelleted by centrifuging at 10,000xg for 15 min at 4°C. The supernatant was transferred to a 1.5 mL tube and KCL was added to 150mM. The sample for input could be taken out for RNA purification or western blot if necessary. 250 µL IgG Sepharose beads (GE Healthcare) were washed three times with wash buffer (20mM Tris-HCL pH7.5, 5mM MgCl_2_, 150mM KCl,0.1% IGEPAL GA-630, 1mM DTT, 20U/mL Rnase inhibitor). The speed of centrifuging of the beads was 900g for 3min. The cell lysate was added to the beads, and the mixture was incubated with gentle rotation for 2 hours at 4°C.The beads were centrifuged as before and the 250 µL unbound or flow-through sample was retained for RNA purification and Western blotting. The beads with bound protein and RNA were transferred to a column, washed with wash buffer 3 times (10mL each), then transferred into a 1.5 mL tube. The beads were suspended in 500 µL wash buffer, 5 µL TEV protease was added, and the mixture was incubated at 16°C with gentle rotation for 2h. After the reaction, beads were centrifuged down and supernatant containing the protein-RNA complexes was retained; 500 µL wash buffer was added to the beads to harvest remaining released RNA and protein. The bound and unbound fractions were then subjected to RNA purification and Western blotting. Before RNA purification, the sample was treated with proteinase K, and Trifast(Peqlab) was used for RNA purification according to manufacturers’ instructions.

Before RNA sequencing, ribosomal RNA was depleted from the flow-through samples by incubation with specific oligonucleotides and RNaseH (Minia *et al.*, 2016). RNASeq analysis was performed on eluted (bound) and flow-through (unbound) samples, and analysed as described previously (Droll *et al.*, 2013, Mugo & Clayton, 2017, Mulindwa *et al.*, 2018, Leiss *et al.*, 2016). The same analysis method was used for RNA obtained after *ZC3H20* and *ZC3H20+21* RNAi. Briefly, after trimming of primers, the reads were aligned with the TREU927 genome using Bowtie2, allowing 20 alignments per read. Reads that mapped to open reading frames were counted (Supplementary Tables S1 and S2, sheet 4, and supplementary Table S3 and S4, sheet 3). For most subsequent analyses, we used a list of unique genes (modified from (Siegel *et al.*, 2010)), in order to avoid giving undue weight to repeated genes and multi-gene families. Allowing 20 alignments means that with the unique set, the normalised read counts reflect the relative abundance of each mRNA. For pull-downs (Supplementary Tables S1 and S2), reads per million reads were counted and the eluate/flow-though ratios were calculated. An mRNA was classified as “bound” if both ratios were higher than 2. Note that these are not the real ratios because we did not normalise the values according to the amounts of mRNA in the different fractions. Differential gene expression after RNAi was assessed DeSeqU1 (Leiss & Clayton, 2016), a custom version of DeSeq2 (Love *et al.*, 2014).

## Microscopy

To examine cell morphology, dry smears were made and fixed with methanol for 1 min at room temperature, then DAPI-stained. Randomly chosen images were quantified, blind, by an independent observer.

## Data availability

The RNASeq raw data are available at Array Express with the following accession numbers: *ZC3H20*

RNAi: E-MATB-7995; *ZC3H20&21* RNAi: E-MATB-7996; ZC3H20 and ZC3H21 RNA pull-downs: E-MATB-7998. Additional raw data are available from BL or CC on request.

## Acknowledgements

We thank Claudia Hartmann and Ute Leibfried for technical assistance and Kathrin Bajak for designing the oligonucleotides and making some intermediate constructs for pHD2910 and pHD2911, and giving initial assistance with the RIP protocol. Tania Bishola did the PSSA2 RIP-PCR experiment. We also thank Keith Matthews for antibody to PAD1.

BL was supported by a fellowship from the Humboldt Foundation and also by a grant from the Deutsche Forschungsgemeinschaft, Cl112/28-1. Consumables and services were paid from the Deutsche Forschungsgemeinschaft grant and by core funding from the State of Baden Württemberg.

## Author contributions

BL did all of the experiments; KM designed differentiation experiments, helped with bioinformatics and did the unique-copy sequence alignments. CC supervised the work. BL and CC designed experiments, interpreted the results, and wrote the paper.

## Supporting information

**<u>S1 Table</u>**

mRNAs bound by ZC3H20. For details see Key and Legend sheets.

**<u>S2 Table</u>**

mRNAs bound by ZC3H21. For details see Key and Legend sheets.

**<u>S3 Table</u>**

Effect of *ZC3H20* RNAi on the transcriptomes of differentiating EATRO1125 trypanosomes

**<u>S4 Table</u>**

Effect of RNAi targeting *ZC3H20* and *ZC3H21* on the transcriptomes of procyclic Lister427 trypanosomes

**<u>S5 Table</u>**

Simplified summary of the RNA pull-down and RNAi results

**<u>S6 Table</u>**

Plasmids and oligonucleotides

**<u>Supplementary Figure S1</u>**

Alignment of kinetoplastid ZC3H20, ZC3H21, and ZC3H22 homologues. The alignment was created using Clustal Omega (https://www.ebi.ac.uk/Tools/msa/clustalo/) (Sievers *et al.*, 2011) with default settings. Only the region containing two zinc finger domains is shown. Organism abbreviations are: CFAC: *Crithidia fasciculata*; Lmj: *Leishmania major*; EMOLV: *Endotrypanum monterogeii*; PCON: *Paratrypanosoma confusum*; TRSC: *Trypanosoma rangeli*; TCCLB: *Trypanosoma cruzi*; Tb: *Trypanosoma brucei brucei*; TCIL: *Trypanosoma congolense*; Tv: *Trypanosoma vivax*. The text colour code is blue - negative charge; purple - positive charge; green - polar; red - non-polar. Underlying colours are as in Fig 1C.

**<u>Supplementary Figure S2</u>**

Alignment of CCCH domains from *T. brucei* proteins that have two CCCH domains.

The alignment was created using Clustal Omega (https://www.ebi.ac.uk/Tools/msa/clustalo/) (Sievers *et al.*, 2011) with default settings. Only the zinc finger domains, with the surrounding 8 residues, were included. “a” is the first (N-terminal) motif, “b” is the second motif. The cladogram with real branch lengths is on the left of the alignment. The text colour code is as in Supplementary Fig S1. The pink area is the domain itself, while the blue and green areas highlight the immediate context which may also affect binding specificity.

**<u>Supplementary Figure S3</u>**

Genotypes of cell lines

A. Maps showing hybridization positions of oligonucleotides used to verify the genotypes of transgenic trypanosomes.

All other panels illustrate genotypes, except panels E and G, which illustrate verification of genotypes C/D and F, respectively, by PCR using genomic DNA as template and oligonucleotides shown in (A). Other lines were verified similarly.

**<u>Supplementary Figure S4</u>**

Additional results concerning developmental regulation

A. TAP-ZC3H20 expression is induced in pleomorphic bloodstream forms at high density. EATRO1125 bloodstream forms expressing N-terminally TAP-tagged ZC3H20 (Z20) were used (Fig S3B). Initial growth (left panel) was in HMI9 at 37°C with methylcellulose. At time point c, cis-aconitate was added to trigger differentiation and the temperature was lowered to 27°C. 1 day later the cells were transferred to MEM-Pros medium (right panel). Cells were harvested at different time points (a-i) for Western blot analysis. Left panels shows cell densities (left) or cumulative cell numbers (right) and the right-hand panel show the results of Western blotting. Ribosomal protein S9 serves as a loading control.

B. TAPZC3H20 expression is induced in monomorphic bloodstream forms at high density. The experiment is similar to that in (A) except that monomorphic Lister 427 bloodstream forms were used. These cannot make stumpy forms and grow to a higher maximum density than the EATRO1125 cells. The parasites were cultivated either in the presence (+MC) or the absence (-MC) of methyl cellulose. The upper Western blot shows TAP-ZC3H20 expression. The lower blot shows typical results for exponentially-growing, independently-generated, bloodstream-form (B) and procyclic-form (P) TAP-ZC3H20 lines. The difference in signal intensity was 3.5×.

C. Expression of ZC3H20 at high density does not require methyl cellulose. EATRO1125 bloodstream forms expressing N-terminally TAP-tagged ZC3H20 were grown to high density in the absence of methyl cellulose and expression of TAP-ZC3H20 was assayed.

D. ZC3H21 expression (Z21) is induced in pleomorphic bloodstream forms upon cis-aconitate (CA) addition. The experiment was like that in panel (A), except that cells expressing TAP-ZC3H21 were used.

E. TAP-ZC3H20 expression increases with a temperature shift to 27°C, but addition not as much as when cis-aconitate is included.

F. TAP-ZC3H20 expression does not increase when Lister 427 procyclic forms are grown to high density.

G. TAP-ZC3H20 expression increases when bloodstream forms are incubated at 19°C for 6h.

H. TAP-ZC3H21 is increased when bloodstream forms are incubated at 19°C for 3-6h. Cell numbers are on the left.

**<u>Supplementary Figure S5</u>**

Ectopic expression of ZC3H20-myc restores EP expression but inhibits growth.

A. Repeat of Fig 5A. EATRO1125 *T. brucei* were used in which lacked *ZC3H20* genes (HKO) but had a tetracycline-inducible copy of ZC3H20-myc. Two clones, C1 and C2, were compared and wild-type was used as a control. At time point 0, long slender bloodstream-form cells were treated with cis-aconitate with or without tetracycline. Cumulative cell numbers are on the left and protein expression on the right.

B. TAP-ZC3H20 is detectable in procyclic forms 6h after protein synthesis inhibition. Cells were treated with 100µg/ml cycloheximide and the TAP tag was detected by Western blotting.

C. Procyclic forms with tetracycline-inducible ZC3H20-myc (Cell line S3 J) were incubated with two different concentrations of tetracycline. Cells were diluted to 1 × 10^6^/ml one day later. Growth is on the left, the Western blot on the right. Lethality of extra ZC3H20-myc was also seen in the presence of wild-type *ZC3H20* genes (not shown).

**<u>Supplementary Figure S6</u>**

Ectopic expression of ZC3H21-myc does not complement the lack of ZC3H20 in differentiating bloodstream forms.

A. EATRO1125 bloodstream forms without ZC3H20 (homozygous knock-out, HKO) were complemented with inducible ZC3H20-myc or ZC3H21-myc. Differentiation of cells was induced at time 0, at 1×10^6^/ml. Cell densities are on the left and a Western blot on the right. Altered EP procyclin expression after 2 days’ ZC3H20-myc induction was seen in one culture (double arrowhead), but not the other (single arrowheads)

B. Morphologies of cells at the outset and 4 days later (also 8 days for wuild-type). The images are differential interference contrast with DNA stain in dark blue.

**<u>Supplementary Figure S7</u>**

RNA immunoprecipitation results

A. Principal component analysis for both experiments. EL= eluate (bound RNA); FT = flow-through (unbound RNA).

B-E. Correlations between replicate experiments (A and B). Each dot represents a single unique open reading frame. Reads per million reads are plotted on a log_2_ scale.

F-I. Relationships between bound and unbound fractions. Orange dots are RNAs that were 2-fold enriched in the eluate in both fractions; the blue dots in H and I are mRNAs encoding flagellar proteins. Results for ZC3H20 and 21 and a few additional outliers are not included.

**<u>Supplementary Figure S8</u>**

Relationship between RNA binding and developmental regulation or mRNA length

A: The x-axis shows the average ratio of bound to unbound normalised read counts for ZC3H20, on a log_2_ scale (Supplementary Table S1). The y-axis shows the ribosome footprint counts for procyclic forms, divided by those for bloodstream forms. The genes showing the strongest mRNA binding are labelled.

Gene numbers are truncated for clarity: “Tb927.” has been removed. Each dot represents a single unique open reading frame.

B. As (A), but for ZC3H21 (Supplementary Table S2)

C: The x-axis shows mRNA lengths on a log_2_ scale. The y-axis has the average ratio of bound to unbound normalised read counts for ZC3H20, (Supplementary Table S1). Coloured dots are in the classes cytoskeleton (orange), ribosome (cyan) and membrane (violet).

D. As (C), but for ZC3H21 (Supplementary Table S2)

**<u>Supplementary Figure S9</u>**

Bound/unbound ratios for mRNAs encoding proteins of different functional classes.

Class assignments are in Supplementary Tables S1 and S2. The average ratio of bound / unbound RPMs for each mRNA were included. The thick horizontal line is the median, and boxes extend to the 25th and 75th percentiles. Whiskers extend to the furthest measurement that is within 1.5× the inter-quartile distance. Other spots are outliers. The grey background box indicates the inter-quartile range for the whole dataset. Coloured boxes are those mentioned in the text, and others for which the median is outside the inter-quartile range.

**<u>Supplementary Figure S10</u>**

ZC3H20 binding to gene family mRNAs.

Each read was assigned once to the best possible location and any reads that aligned twice were excluded (MAPQ score ≥ 42), such that only unique reads were detected. The coverage was visualised and screen shots are shown.

A. Specific pull-down of *ZC3H20*, but not *ZC3H21* or *ZC3H22*. The 5’ ends of the *ZC3H20* and *ZC3H21* coding regions have very few mapped reads because they were eliminated by the mapping criteria.

B. Pull-down or *EP1* and *GPEET* procyclin mRNAs, but not *PAD1* or other mRNAs. The lack of reads mapping to *EP2* and *EP3* might be caused by sequence identity.

C. ZC3H20 binds to *PSSA2* and *MCP12*, but not *PTP1* mRNA. (Enrichment of *TRF* is lower than *MCP12*, less than 1.45×)

D. Reads mapping to trans sialidase mRNAs, Tb927.7.6850 has no reads because this sequence is identical in other trans sialidase genes.

**<u>Supplementary Figure S11</u>**

ZC3H21 binding to gene family mRNAs.

Each read was assigned once to the best possible location and any reads that aligned twice were excluded (MAPQ score ≥ 42), such that only unique reads were detected. The coverage was visualised and screen shots are shown.

A. Specific pull-down of ZC3H21, but not *ZC3H20*; also mMDH and mRNA encoding a pteridine transporter. The 5’ ends of the *ZC3H20* and *ZC3H21* coding regions have very few mapped reads because they were eliminated by the mapping criteria.

B. Specificity for PAD2, but not PAD1; and pull-down or *EP1* and *GPEET* procyclin mRNAs. The lack of reads mapping to *EP2* and *EP3* might be caused by sequence identity. Two different scales are shown.

C. ZC3H21 does not bind to unique sequences in trans sialidase mRNAs. (BInding to repeated sequences cannot be assessed from this alignment).

**<u>Supplementary Figure S12</u>**

Effects of RNAi targeting ZC3H20 and ZC3H21 simultaneously in procyclic forms.

Effects on mRNAs encoding proteins of different functional classes are shown. Class assignments are in Supplementary Tables S1 and S2. The effects of RNAi on each mRNA (log_2_ fold change from DeSeq2) were included. The thick horizontal line is the median, and boxes extend to the 25th and 75th percentiles. Whiskers extend to the furthest measurement that is within 1.5× the inter-quartile distance. Other spots are outliers. The grey background box indicates the inter-quartile range for the whole dataset. Coloured boxes are those mentioned in the text, and others for which the median is outside the inter-quartile range. Selected outlier spots with decreased expression are labelled, and of these, those that were significantly reduced (Padj <0.05) are filled. AT: adenosine transporter; FH: fumarate hydratase; PEPCK: phospoenolpyruvate carboxykinase; PPDK: pyruvate phosphate dikinase; MDH: malate dehydrogenase; TS: trans-sialidase; GCS: gamma glutamyl cysteine synthase.

**<u>Supplementary Figure S13</u>**

Relationship between ZC3H20/21 RNAi and RBP10 expression in procyclic forms

The effects of RNAi targeting ZC3H20 and ZC3H21 simultaneously are on the y-axis and the effect of expressing RBP10 (Mugo & Clayton, 2017) on the y-axis.

