## Supplementary material for "The zinc finger proteins ZC3H20 and ZC3H21 stabilise mRNAs encoding membrane proteins and mitochondrial proteins in insect-form *Trypanosoma brucei*": Fig S1

```
TRSC58_03512-t26_1     -------MVVVDPATRKLLIPCNCIYPTRAQ-QRAAIPSLCQLFLQGRCRQGAQCHQVHA   52
TCCLB.506859.204       DADGTLCMVVVDPATRKLLIPCNFIYATRAQ-HRATIPSLCQLFLHGRCRQGIQCHQVHA   75
CFAC1_240014600.1      NSDGELCMTVVDPVTRKLLIPTQYIFDTRAQ-QRGTLPSLCQLFLLGRCRQGQGCYQVHA   73
LmjF.22.0740           NADGELCMTVVDPVTRKLLIPTQYIFETRAQ-QRGTLPSLCQLFLSGRCRKGQGCYQLHA   73
EMOLV88_220011900.1    NADGEVCMAVVDPATRKLLIPTRYIFPTRAQ-HRGTLPSLCQLFLAGRCRKDQGCYQVHA   73
PCON_0075640           GSDGIARMVTVDPLTRKLHIPCEFVFPTRAS-QRGTLPSLCQLFLNGRCRLGSNCYQVHA  193
Tb927.7.2670           DEKGVLCLTVVDPATRKLLIPLSCVYPTQAQ-QRTTIPSLCQLFLNGRCRQGTQCHQVHA   83
Tb927.7.2660           FEKGVECMVVVDPATRKLRIPLSCVYPTRAH-QRTTIPSLCLLFLDGRCRQGTQCHQVHA   75
TCIL3000_7_1980.1      DDKGVLCMTVVDPVTRKLLIPLNHVYPTRAQ-QRTTIPSLCQLFLNGRCRQGAQCHQVHA   75
TvY486_0702540         CHDGQLCMTVVDPATRKLLIPLEYIYVTRAQ-QRSTIPSLCQLFLQGRCRQGAQCHQVHA  159
TRSC58_00606-t26_1     -------MVVVDPATRKLLIPCNCIYPTRAQ-QRAAIPSLCQLFLQGRCRQGAQCHQVHA   52
TCCLB.506859.230       RADGTLCMVVVDPATRKLLIPCNFIYATRAQ-HRATIPSLCQLFLHGRCRQGTQCHQVHA   83
                              ***    ** :*    :  * *    *   ***C  *:    *C     C    H:
TCCLB.511817.20        KCGEEKVIVVVDPRRSKLRVPLSAISPTKALGIRAQNPSLCLLFQSGRCRQGVNCHQVHV  239
DQ04_01931010-t26_1    KEGKKETVPVVDPQRAKLHVPLSAITPTKALGGRVQHPSLCLLFQSGRCRQGANCYQMHV  313
TvY486_0702570         AVEGERKIPVVDPQRSKLHVPFSAIFPTKALGGRVSFPSLCLLFQAGRCLRGSNCYQMHI  294
TCIL3000_7_2020.1      DPEEEQRIPVVDPQRSKLHVPLSSIIPTKALGGRVSFPSLCLLYQSGRCFRGLAVIKCTL  310
Tb927.7.2680           QPTEEQRIPVVDPQRSKLHVPLSSIIPTKALGGRVSFPSLCLLFQSGRCLRGASCYQMHI  312
CFAC1_240014700.1      NVDERGCITVVDPQRRKLHVPLSAIQTTKALNGRLKTPSLCLLFQSGRCRQGDNCYQVHV  618
LmjF.22.0750           RIDDRGCIAVVDPQRRKLHVPLSAIHMTKALNGRLKTPSLCLLYQLGRCRQGENCYQVHV  740
EMOLV88_220012000.1    RIDGGGCVTVVDPQRRKLRVPLSAIQMTKALNGRLKTPSLCLLYQSNRCRQGESCYQVHV  729

DQ04_15331000-t26_1    -------------------------------MEYGSCLA---------------------    8
TRSC58_03512-t26_1     NVDVVMVLRDQVGNLPRCCPFHGDDDIAGVLNERSLLS----------------------   90
TCCLB.506859.204       SLDAVVALRSRVGKLPCCCVFHGDEDIADVLNERSWLS----------------------  113
CFAC1_240014600.1      DWEAVQRLRAQVDSLPCCCPGHGDKDHLGIAENVPLRDVLSGNGGH--HHGNTHASAAEA  131
LmjF.22.0740           DWEAVQRLRSQVDSLPCCCPTHGDKDHIGVLENALLRESVSLGHPHSHHHGNSEQNSADA  133
EMOLV88_220011900.1    ECEAVRYLRSQVDSLPCCCHTHGDKDHMGVLENAPLWNFFSQNQAHAPGDARSHQCSVDA  133
PCON_0075640           DVHVVAQLRAQVDSLPRCCAFHGDKDHVHSLHH-PLRD----------------------  230
Tb927.7.2670           ALDVVAALRSQVDYLPICCALHGDRDYVNALDNRSWMS----------------------  121
Tb927.7.2660           ALNVVAALRSQVDYLPICCALHGDRDYVNALDNRSWMS----------------------  113
TCIL3000_7_1980.1      SPDMVTVLRSQVETLPRCCALHGDRDYVGVMDDQSWVS----------------------  113
TvY486_0702540         PLEIVSALRSQVETLPWCCAEHGDRDYAGVLNEQSWVS----------------------  197
TRSC58_00606-t26_1     NVDVVMVLRGQVGNLPRCCLFHGDDDIAGVLNERSLLS----------------------   90
TCCLB.506859.230       SLDAVVALRSRVGKLPCCCVFHGDEDIADVLNERSWLS----------------------  121
                           *   **        *  **   ***
TCCLB.511817.20        DPEIVQRLRRIVDSLPCCCTFHGDCN-------THCWD---------------------A  271
DQ04_01931010-t26_1    DPEVVKELRQIIQGLPCCCTYHGDCN-------SKLWD---------------------A  345
TvY486_0702570         DPDVVRRLRQINESLPFCCTFHGDCN-------SKMWD---------------------A  326
TCIL3000_7_2020.1      SPE
Tb927.7.2680           DPQVVQRLRQINESLPYCCAFHGECN-------ANKWD---------------------A  344
CFAC1_240014700.1      DPATVDRLRIDVENMPCCCLLHGDCN-------CHLVD---------------------P  650
LmjF.22.0750           DSAIVERLRADAKNTPCCCFQHGDSN-------GQVMD---------------------R  772
EMOLV88_220012000.1    DPATVERLRTEARDMPCCCFHHGDGN-------SHLMD---------------------L  761

DQ04_15331000-t26_1    -----------NVWIHIPESSFNG-GFIPLSRTGYTIPVAKMLNELPQSIHMELFRYRQ-   55
TRSC58_03512-t26_1     -----------KVVLYIPEVSFEG-GYVPLSRIGYTVPISKMLNKRDSDVVRALDSYRQ-  137
TCCLB.506859.204       -----------KVVLYVPEVSFEG-GYIPLSRVGYTVPISKILNEVSHEIAHSLESYTE-  160
CFAC1_240014600.1      AAVAGSTVCARDVVVYVPGCSTYQGSYVPLERVSYTIGLRRLLEEQQVLLAAPTAHVSL-  190
LmjF.22.0740           AAAAESTVGANDVVLYVPGCSFFEGSYAPLDRVSYTVGLRRLLEEQRVLLTPSTARISL-  192
EMOLV88_220011900.1    TAAPQSTIHTNDIVIYVPGCSFFEGGYVPLVRVSYTNGLRRLLEEQCVLFPPSTAQVTL-  192
PCON_0075640           -----------RLFVKLPGSTWGGDGNIPLHRLSFTTALRWLLGENKDVIAALHQGLPKL  279
Tb927.7.2670           -----------RVVVHVPDATYGG-GYIPLARFSYTTPISRILREVNARL----------  159
Tb927.7.2660           -----------RVVVHVPDATYGG-GYIPLARFSYTTPISRLLDALFHSTP---------  152
TCIL3000_7_1980.1      -----------RVVVHVADAAYEG-GYIPLSRFSYTIPVSKMLNRLPLSAI---------  152
TvY486_0702540         -----------RVVVHIADVQYEG-GYIPLSRFSYTLPVSKLLGEYGNRLE---------  236
TRSC58_00606-t26_1     -----------KVVLYIPEVSFEG-GYVPLNRFSYTPALRRVLKEKQGRLM---------  129
TCCLB.506859.230       -----------KVVLYVPEVSFEG-GYIPLSRVSYTVALQRILKEKPDRAI---------  160
                                                  :*:     :*   :    :
TCCLB.511817.20        KANSKRGV------------FISN-VWVPLSHVAYTSGLARFV-TNKVQRP---------  308
DQ04_01931010-t26_1    KANADRTI------------VINR-VAVPLVRVAYTSGLSRFV-KNEMQRP---------  382
TvY486_0702570         QGRANHTI------------TIKG-VTVPLSHVAYTNGLERFVLKSNHSGS---------  364
Tb927.7.2680           EANAHRTI------------LIKG-AAVPLSRVAYTNG ERFVLKNNASRS---------  382
CFAC1_240014700.1      ACYEGRSL------------LIAGQYNVPLSRVAYTAGLQRVLQEEAMSVP---------  689
LmjF.22.0750           TAYEGRSL------------GIAGQFSVPLTRVAYTAGLQRVLQDQPACAP---------  811
EMOLV88_220012000.1    TAHAGRSL------------DIVGQLSLPITRTAYTAGLQRVLQDEQPCAI---------  800

DQ04_15331000-t26_1    ------------AVREGRM-PIADPPRVVLNAADIPICRLHIL-DRCRYAEECSFLHLCK  101
TRSC58_03512-t26_1     ------------SLQEGSA-PPEGSPVVVLEAGDIPICRLHVK-ERCRFAEECGFLHLCR  183
TCCLB.506859.204       ------------AIREGRV-PHEESPVVVLEAGDIPICRLHIR-ERCRFAEECSFLHLCK  206
CFAC1_240014600.1      ----------------LNAETGFPEDKIVADASGATVCRLHAM-DRCRYAEECKFLHLCK  233
LmjF.22.0740           ----------------LNQVTGFPEEKIVADASAATVCRLHAM-GRCRYAEECKFLHLCK  235
EMOLV88_220011900.1    ----------------LNAATGTTERKIVADASGATMCRLHAM-DRCRYAEECKFLHLCK  235
PCON_0075640           PRDIDSEGFNVTAVVDSPTGNGRGNSIVLAAGESIAVCRLHIA-DRCRYAEDCKFLHICK  338
Tb927.7.2670           ----------ESGVSVAAVD-GGGHPKRMVLNACDFKICGLHTL-DRCRYAEECIFLHICK  208
Tb927.7.2660           ----QAEG-QDGAQWESRA-FGTPPSCLVLEAGDITLCRLHVQ-DRCRYAEECNYLHICK  205
TCIL3000_7_1980.1      ----RAACYHKQLQQEGRP-HDVASLRVVLEAGDIPLCRLHVL-ERCRYAEECSFLHICK  206
TvY486_0702540         ----PDEGGSGG---GGDS-SGRGKQRVILNALDMKICNLHVF-DRCRYAEECIFLHVCK  287
TRSC58_00606-t26_1     ----PDNGDKEIC--GGSD-LQAEGSCLVLNAGDQSICRLHIF-DRCRYADDCRFLHVCK  181
TCCLB.506859.230       ----YELREESTC--GEND-VRPGNSRLIIDASDQSLCRLHIF-DRCRYAEECRFLHLCK  212
                                                            :C **      C*:    :C  : :H:*
TCCLB.511817.20        ----------------LSF---------------GVLCRLHGKTGGCRYGADCWYLHVCR  337
DQ04_01931010-t26_1    ----------------LST---------------GVLCRLHGAPGGCRYGADCWYVHVCR  411
TvY486_0702570         ----------------LNT---------------CAICRLHCRPGGCRYGADCWYVHICR  393
Tb927.7.2680           ----------------LNT---------------CAVCRLHGKPGGCRYGADCWYVHICR  411
CFAC1_240014700.1      ----------------VNP---------------SVLCRLHGQRGGCRFGADCKFVHVCC  718
LmjF.22.0750           ----------------VKA---------------SVLCRLHGQPGGCRFGADCRFVHVCR  840
EMOLV88_220012000.1    ----------------VNP---------------SVLCRLHGQPGGCRFGADCRFLHICF  829
```
