## Supplementary figures and images for "The zinc finger proteins ZC3H20 and ZC3H21 stabilise mRNAs encoding membrane proteins and mitochondrial proteins in insect-form *Trypanosoma brucei*"

### Fig S2

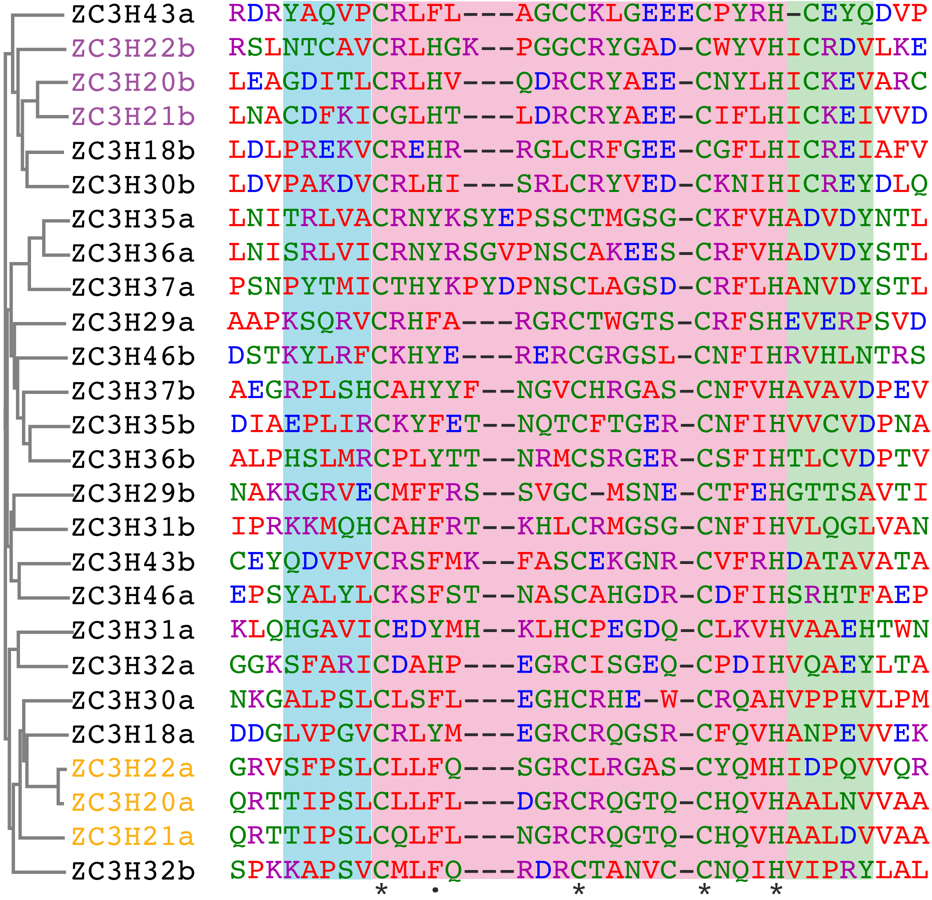

### Fig S3

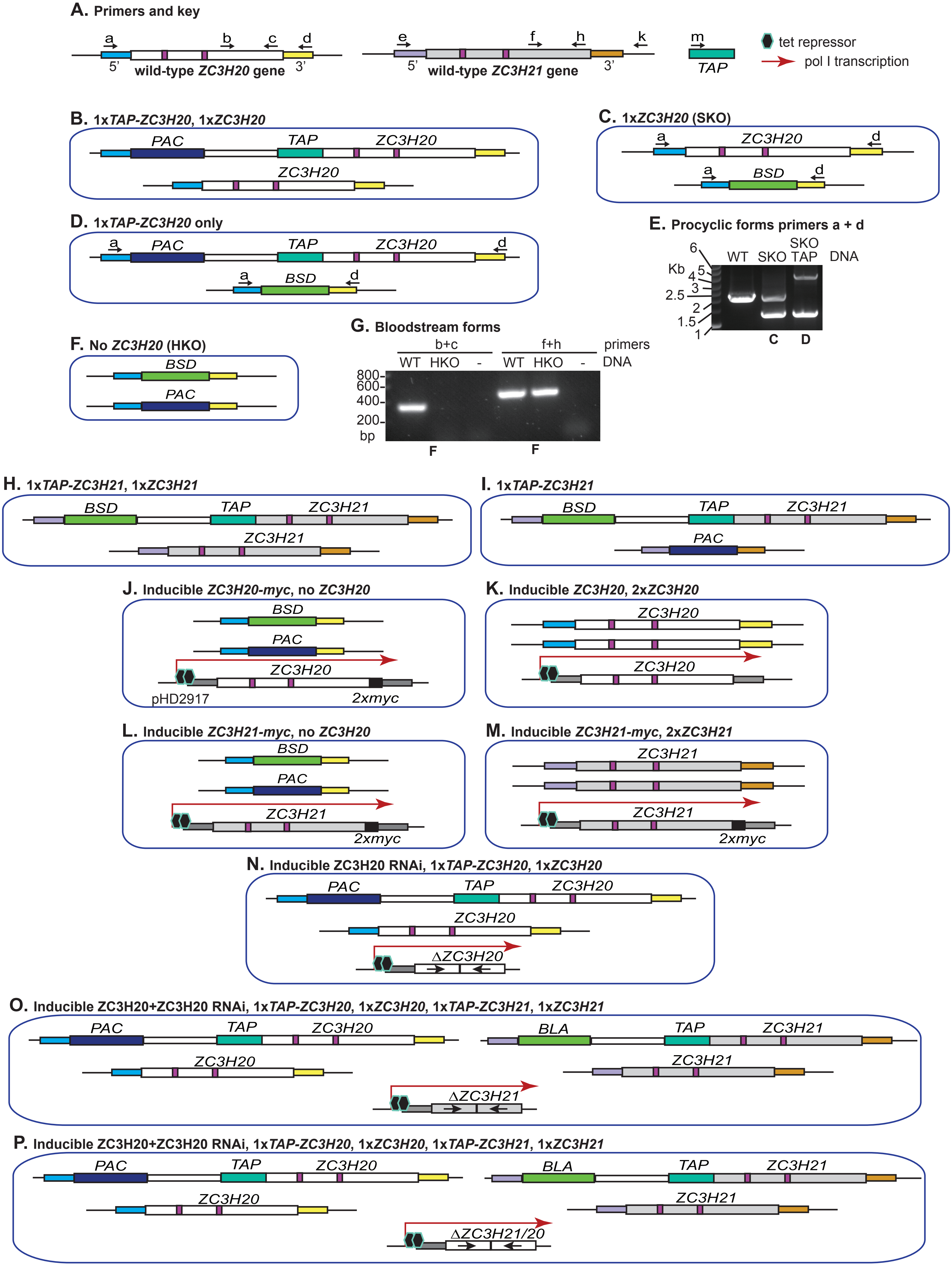

### Fig S4

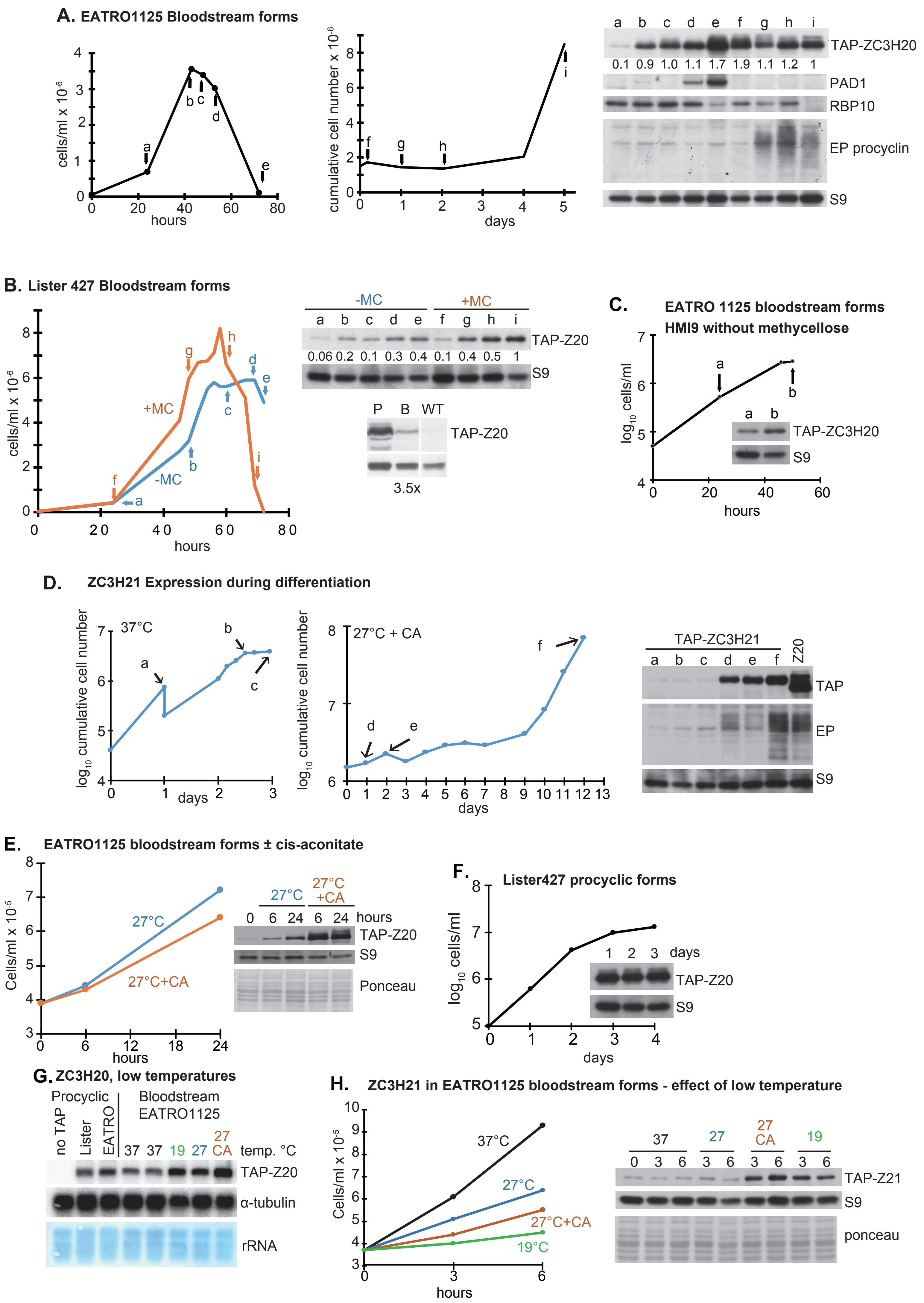

### Fig S6

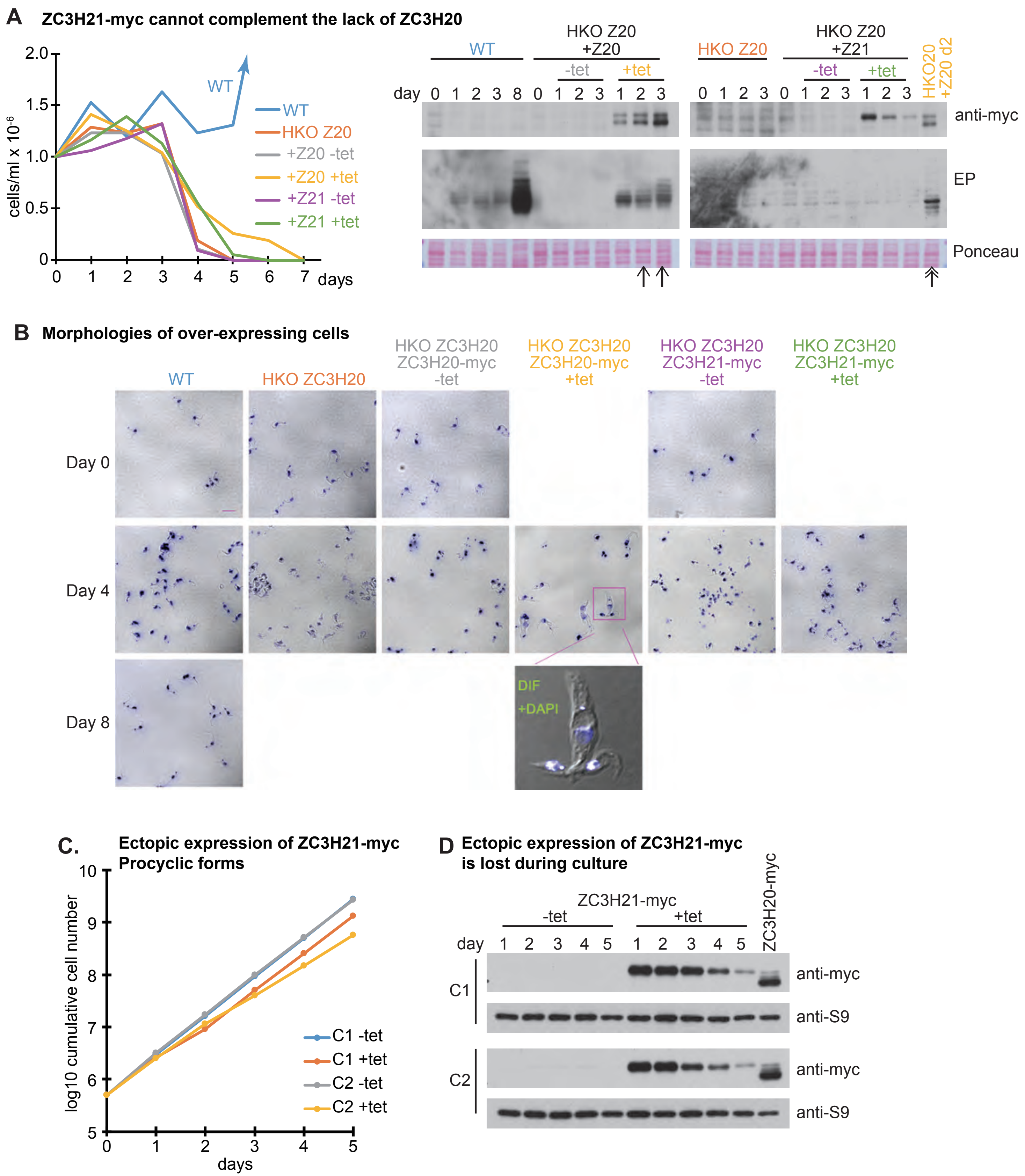

### Fig S7

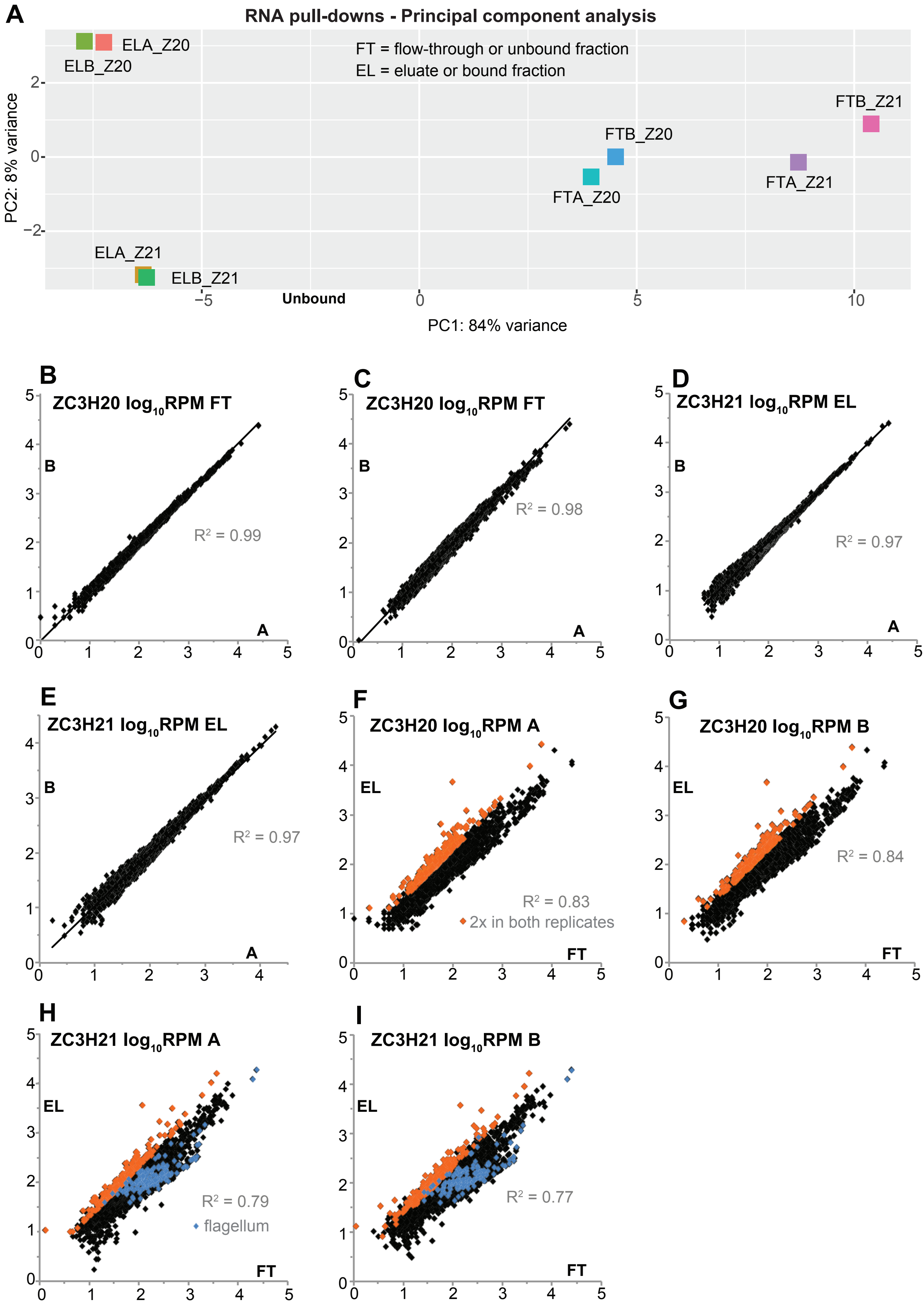

### Fig S8

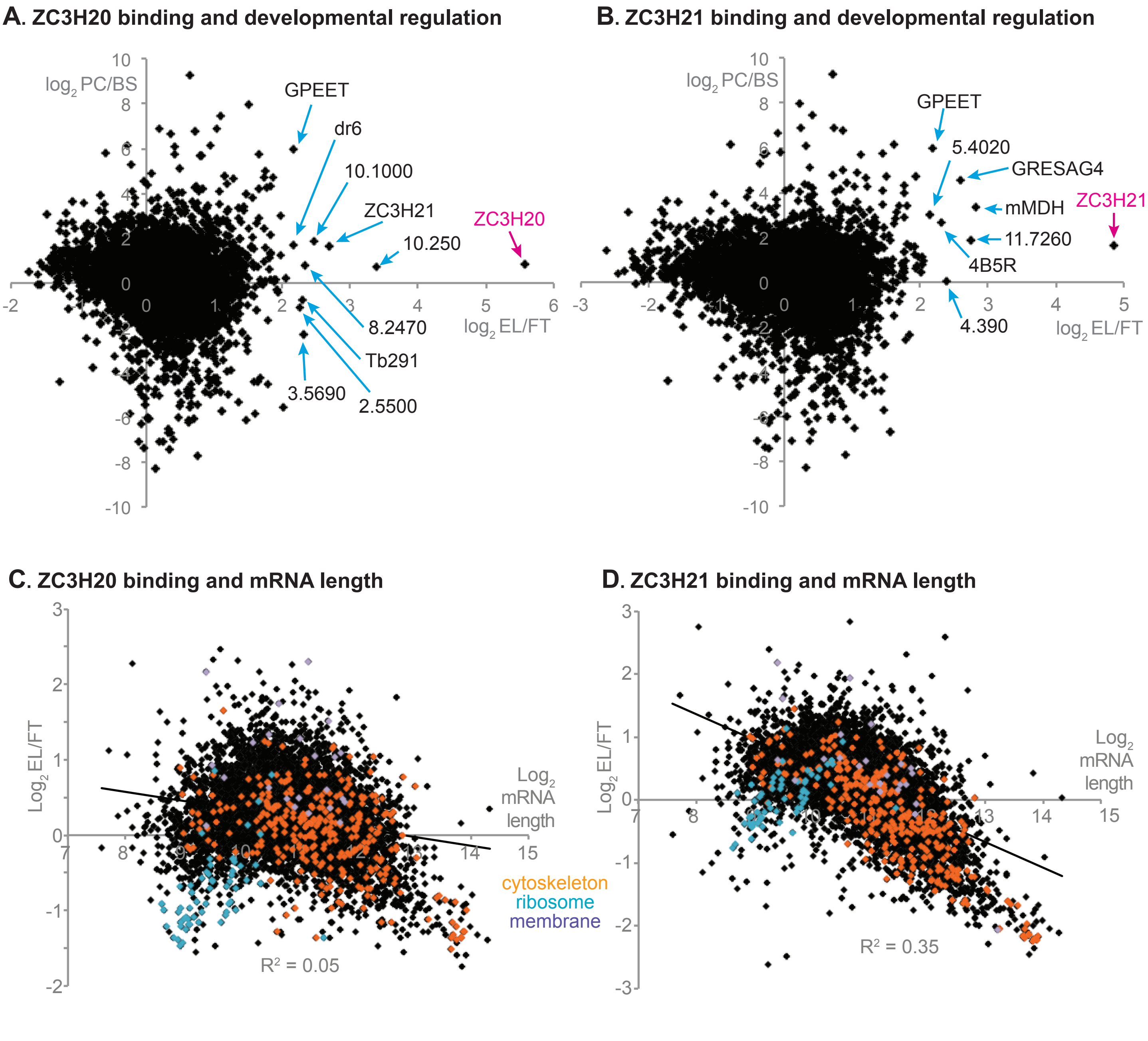

### Fig S9

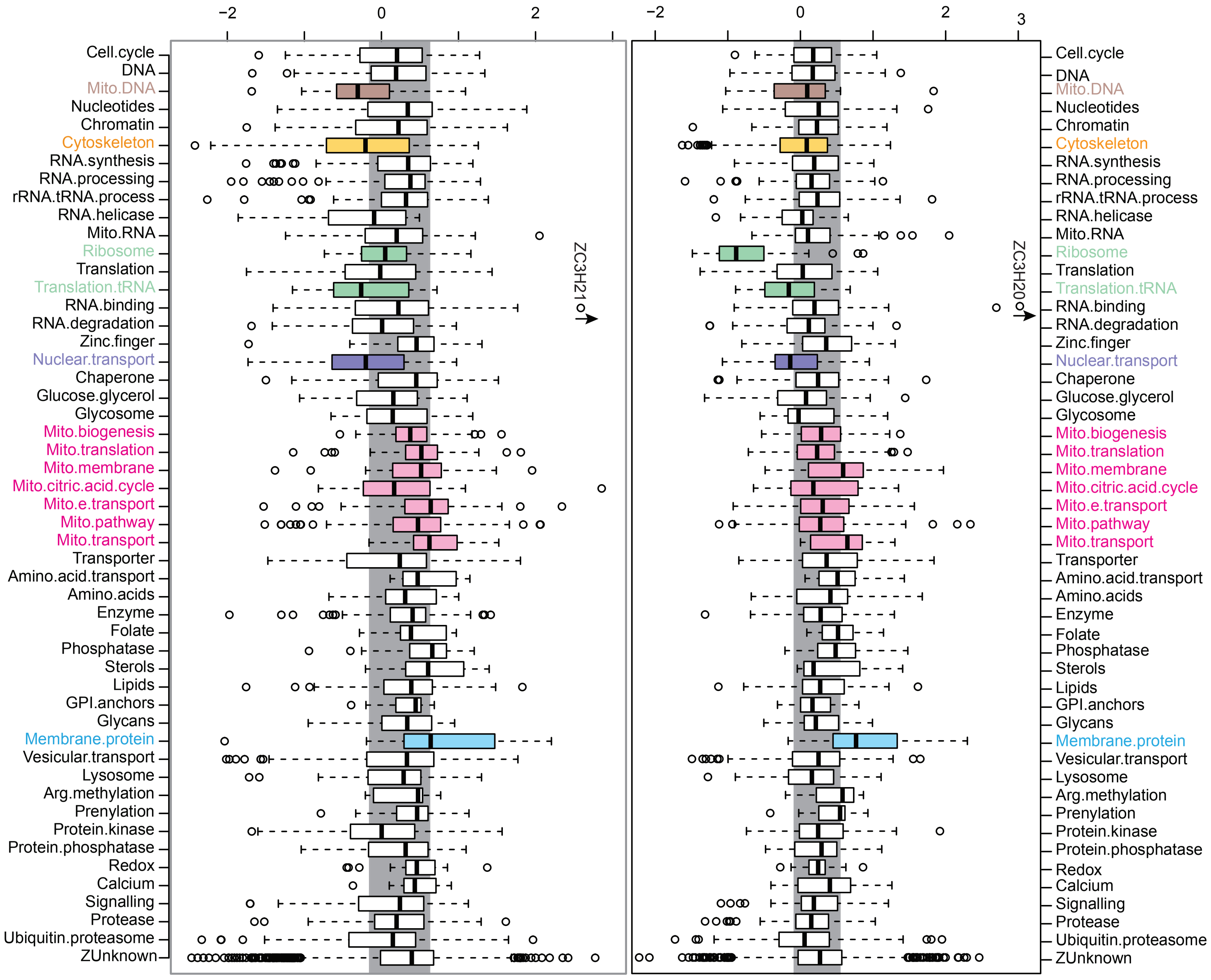

### Fig S10

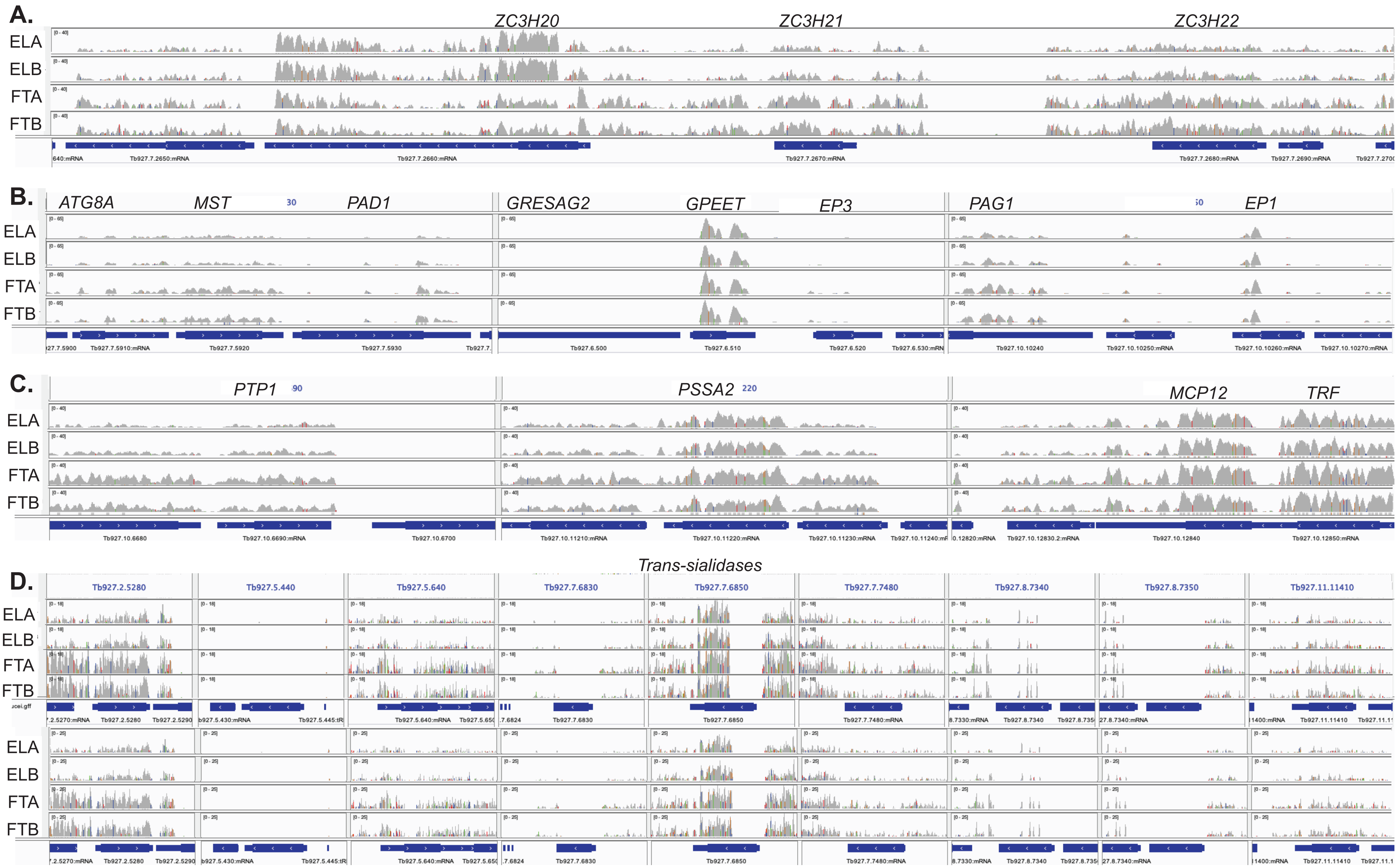

### Fig S11

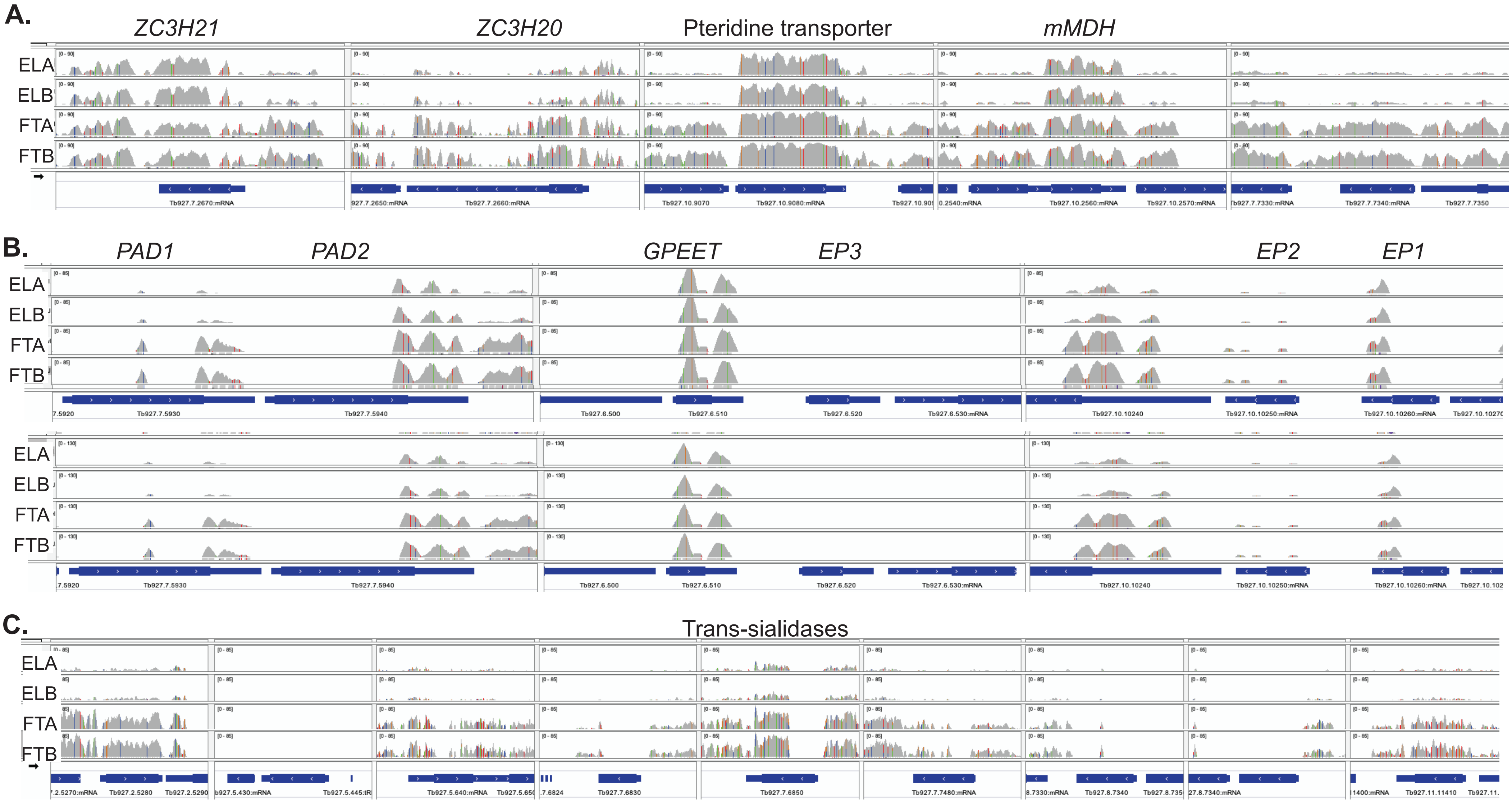

### Fig S12

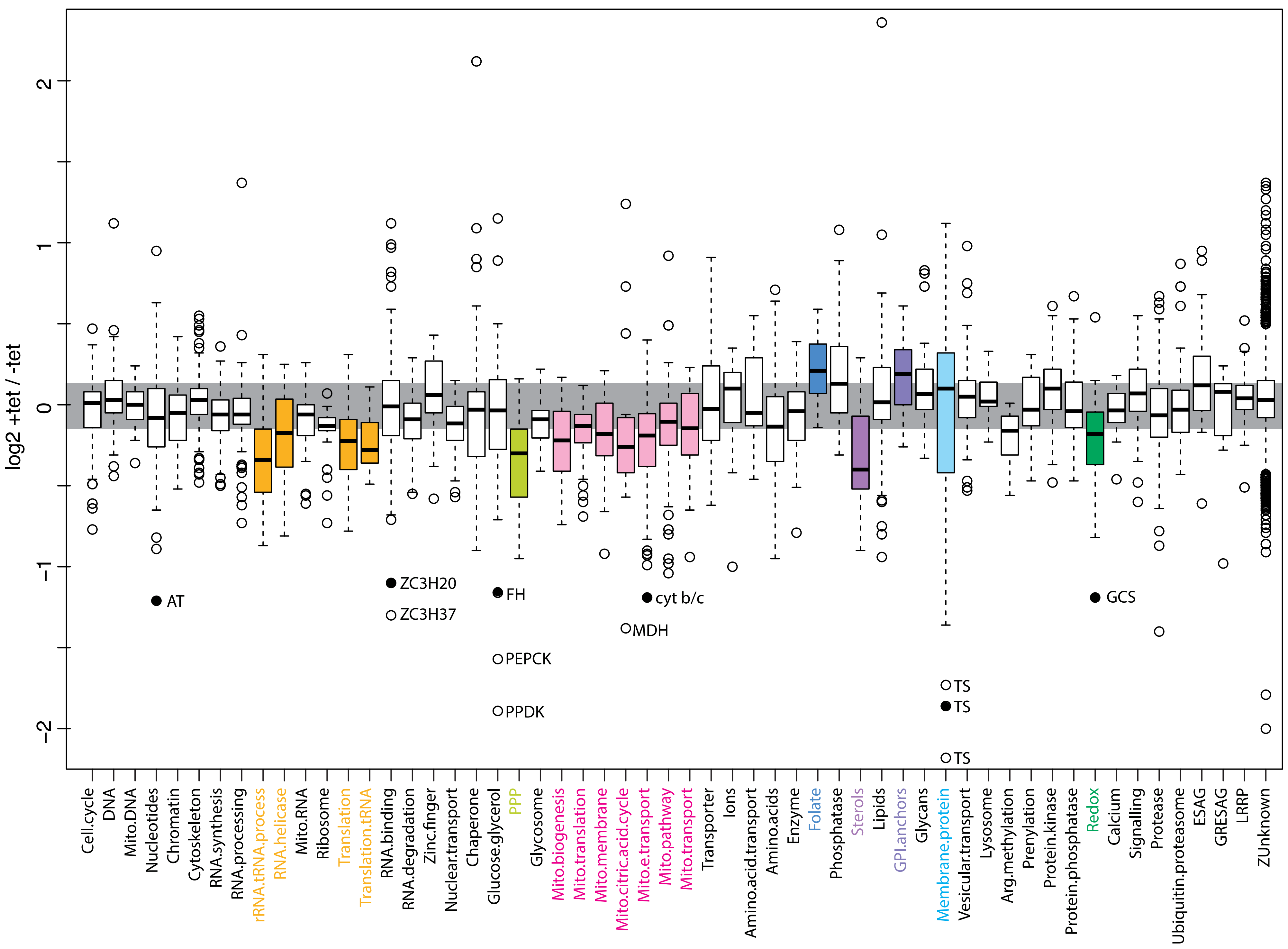

### Fig S13

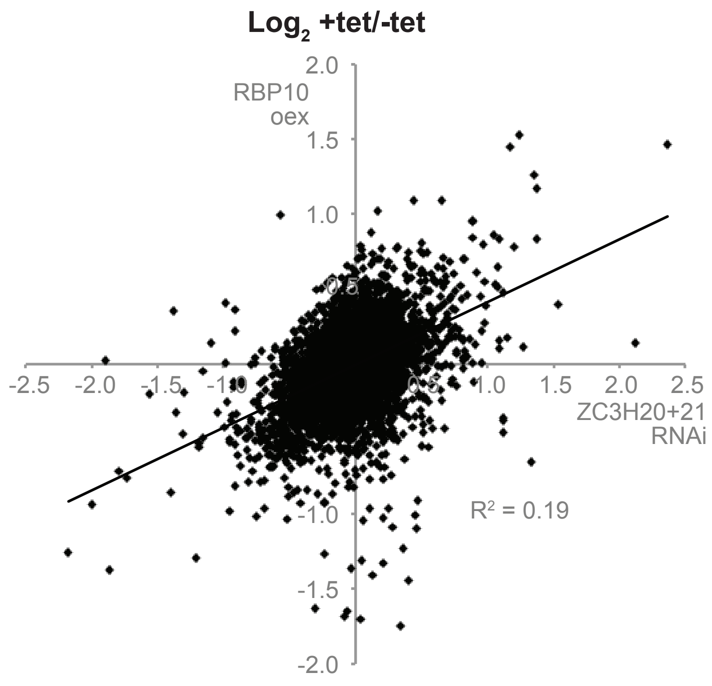
