## Supplementary material for "The zinc finger proteins ZC3H20 and ZC3H21 stabilise mRNAs encoding membrane proteins and mitochondrial proteins in insect-form *Trypanosoma brucei*": Fig S5

**A. Ectopic expression of ZC3H20-myc represses growth after differentiation**

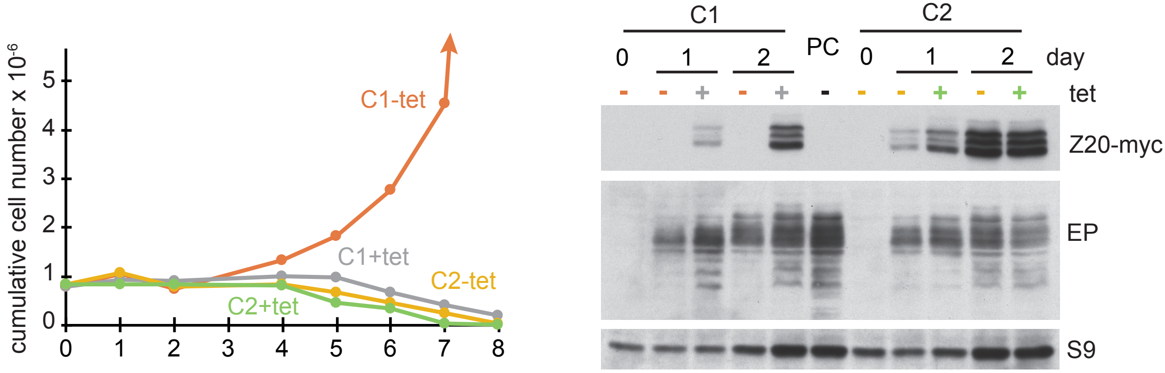


**B. Stability of ZC3H20 in procyclic forms.**

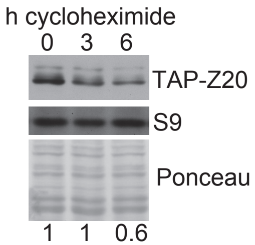


**C. Procyclic forms: Ectopic expression of ZC3H20-myc inhibits growth**

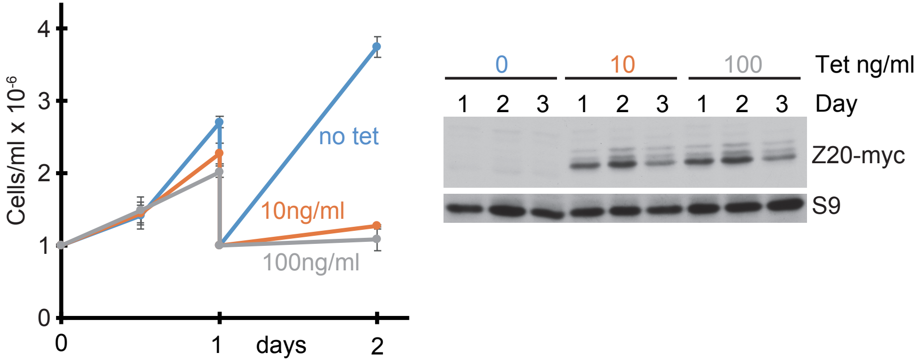
